# Multiparameters dependance of tissue shape maintenance in myoblasts multicellular aggregates: the role of intermediate filaments

**DOI:** 10.1101/2021.12.18.473332

**Authors:** Irène Nagle, Florence Delort, Sylvie Hénon, Claire Wilhelm, Sabrina Batonnet-Pichon, Myriam Reffay

## Abstract

Liquid and elastic behavior of tissues drives their morphology and their response to the environment. They appear as the first insight on tissue mechanics. We explore the role of individual cell properties on spheroids of mouse muscle precursor cells by developing a fully automated surface tension and Young’s modulus measurement system. Flattening multicellular aggregates under magnetic constraint, we show that rigidity and surface tension act as highly sensitive macroscopic reporters closely related to microscopic local tension and effective adhesion. Shedding light on the major contributions of acto-myosin contractility, actin organization and intercellular adhesions, we reveal the role of desmin organization on the macroscopic mechanics of this tissue model.

## Introduction

Tissue-forming cells interact with each other and with their environment [1, 2, 3] giving rise to interesting visco-elastic fluid behaviors [4]. Viscoelastic properties have been observed as determinant both in epithelia [5] and in 3D tissue-forming systems [6]. Physical properties such as surface tension [7, 8] and viscosity [9] can then be introduced to predict tissue organization and shape [10]. Global mechanical characteristics of tissues emerge from individual cell components and their interplay [11, 12, 13]. They thus appear as potential good reporters of individual cell behavior and global organization.

Mechanical characteristics have been measured in minimal tissue models to finely control the initial state and the environment. Indeed multicellular aggregates [14, 9] prove to be powerful tools to apprehend fundamental biological processes [15]. They are able to mimic various biological phenomena [16] but as model systems, are easier to use and to monitor than *in vivo* tissues [17]. Combining simplicity, reproductibility and biological significance, they appear as a system of choice both for biophysics and computation [18, 19, 20].

Myoblasts are highly studied cells to understand myogenesis because of their interest in myopathy modelling [21] and drug testing [22]. Muscle cell mutations implicated in numerous diseases have been extensively studied starting from symptoms to molecular origins identification [23, 24]. However the early effects of mutations in tissue are unclear due to a lack of *in vitro* biomimetic muscle systems [25]. To address these limitations we focus on mouse myoblast cells (C2C12) to test the sensitivity of 3D unorganized early stage muscle tissue models to individual cell modifications. C2C12 are adhesive and highly contractile cells (unsurprisingly regarding their function). Their high assumed surface tension makes them challenging to characterize.

Herein tagging cells with magnetic nanoparticles, thus providing them with magnetic properties while being harmless to cell functioning [26, 27], we design an integrated system to form multi-cellular myoblasts aggregates and to measure their mechanical characteristics. Indeed myoblast spheroids are first molded under various conditions using magnetic nanoparticles tagging [28] then they are flattened under magnetic force to measure their surface tension and Young’s modulus from their profile. Surface tension describes the differences on tissue affinities [29] while Young’s modulus characterizes its elasticity. The interplay between these macroscopic measurements and the molecular or cellular processes represents an appealing way to look at translation of microscopic properties at the tissue-scale. We explore the set-up threshold and demonstrate its efficiency through dose-reponse contractility inhibition. This magnetic tensiometer thus appears as a sensitive system to explore mechanical properties of myoblast-derived tissues. While actin and cadherins role have been implicated in tension surface of multicellular aggregate or embryo [30], the role of intermediate filaments has never been checked. On the contrary to micro-filaments, intermediate filaments are tissue specific. Their function is thus highly related to the specific one of the cells. Disease mutations in intermediate filaments severely impairs individual cell nanomechanical properties [31]. Their network supports the shape of individual cell [32] and withstand applied constrains. Looking at the interplay between their organization and tissue shape maintenance thus appears as primordial. We focus on a type III intermediate filament: the desmin that is specific to muscle cells. We look at cells expressing mutated desmin and exhibiting organization defects of the desmin cellular network to shed light on the crucial role of intermediate filaments on muscle tissue model global mechanics at an early stage of differentiation.

## Results

### Magnetic tensiometer for multicellular aggregates

C2C12 cells are labelled with superparamagnetic nanoparticles without modifying their biological capacities [26] or inducing hypoxia nor apoptosis (Supplementary Figures 1 and 2). It is then possible to organize and stimulate them at will using external magnets. Magnets first drive cells in agarose molds to create spheroids of controlled size [28] and content, as inhibitors or reagents can be added at this stage (Figure 1a). Cohesive spheroids are obtained within 12 hours and imaged (Figure 1b). Their profile is registered while the magnet is approached (Supplementary Figures 3 and 4). Surface tension and magnetic forces in volume compete to determine the equilibrium shape while Young’s modulus is at stake for the contact area [33] (Figure 1c, Supplementary Figure 3). The height, width of the aggregate and the ending points of the contact zone are pointed in the Tensio 𝕏 application (Supplementary Figure 3), a dedicated MatLab interface we developed to fit the obtained profile. The elasticity and the capillary parameter 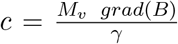 (*M*_*v*_, *γ* and *B* stand for the magnetic moment per unit of volume, the spheroid surface tension and the magnetic field respectively) [14] are extracted. Knowing the magnetic volume moment, the surface tension is easily deduced from *c*. Deformation precision is around 5 *μ*m leading to a precision in the range of 5 *−* 20% for surface tension and 5 *−* 10% for the elasticity of C2C12 multicellular spheroids (error increasing with surface tension as spheroids deform less). This error is smaller than the inherent distribution measured over different aggregates (Figure 2a-b).

**Figure 1:**
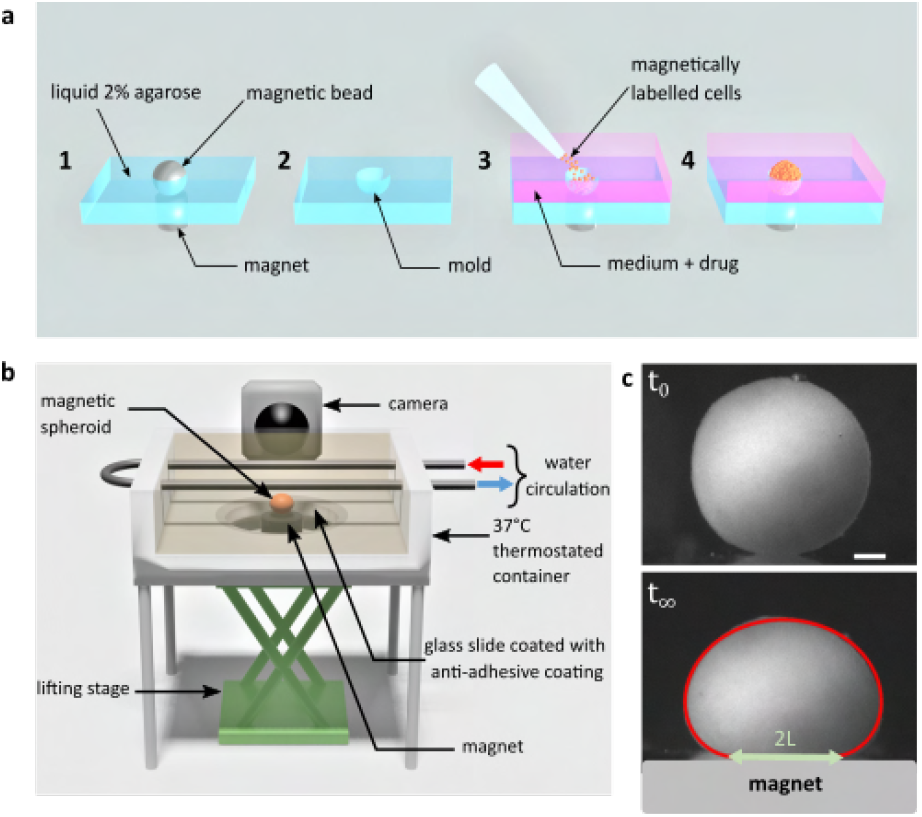
Magnetic tensiometer integrated measurements set-up Magnetic tensiometer set-up. a) Schematic of the magnetic molding process. A network of calibrated size steel beads deposited over cylindrical magnets is embedded in heated 2% liquid agarose (1). After agarose gelling, beads are removed creating a semi-spherical mold (**2**). Magnetically labelled cells are seeded in these non-adhesive treated molds. Magnets placed below each mold drive the cells inside the mold (**3**). Due to high local cell density, cell-cell contacts are created to form cohesive multicellular aggregates within 12 hours (**4**). b) Schematic of the magnetic force tensiometer set-up. A temperature-regulated tank (37 ° C) is sealed with a non-adhesive glass slide to ensure non-wetting conditions for the multicellular spheroid formed by magnetic molding (Methods and Figure 1a). A cylindrical neodymium permanent magnet (6mm diameter and height, 530 mT, grad(*B*) = 170T:m ^− 1^) is positioned underneath and approached thanks to a lifting stage. The tank is filled with transparent culture medium and the aggregate side profile is monitored with a camera. c) Representative pictures of C2C12 spheroid profiles. Top and bottom pictures show respectively a C2C12 spheroid before the magnet approach (t_0_) and under magnetic flattening when the equilibrium shape is reached (t_1_). The dynamics is shown in Supplementary Figure 4. The spheroid surface tension is measured by fitting the aggregate shape with Laplace profile (red line) while the elastic modulus is extracted from the radius of the contact zone L (green arrow) using Hertz theory as described in [28] (Supplementary Figure 3). On this picture γ = 21mN/m and E = 100Pa. Scale bar = 200 *μ*m.

**Figure 2:**
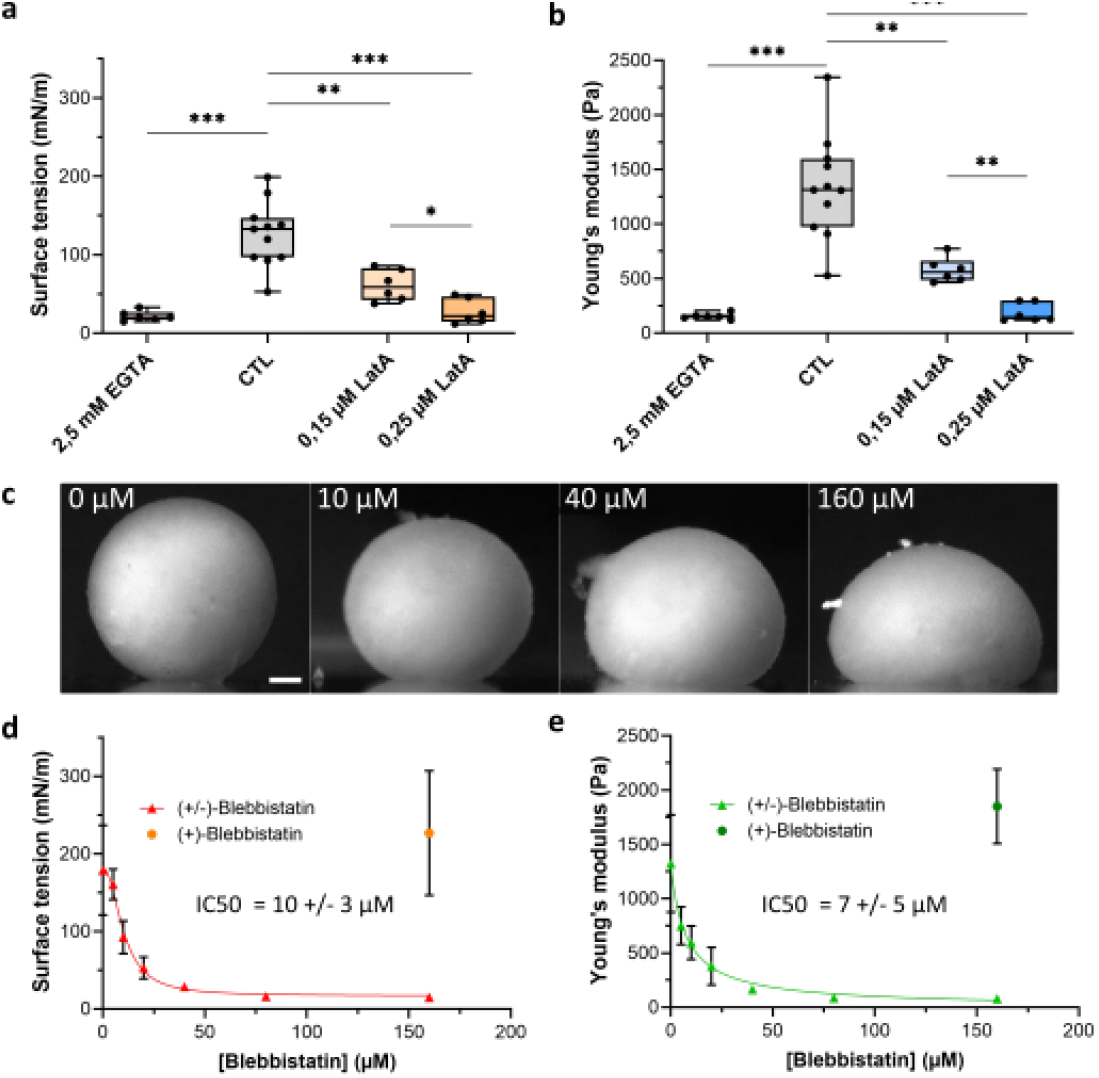
Co-action of cortical tension and intercellular adhesion in multicellular spheroid surface tension and Young’s modulus. **a-b)** Variation of surface tension (a) and Young’s modulus (b) of C2C12 spheroids for 2.5 mM EGTA (calcium chelator), 0.15 *μ*M and 0.25 *μ*M latrunculin A (actin disruptor). Floating bars represent min to max variations and the midline indicates the median. **c)** Representative pictures of C2C12 spheroids under magnetic flattening at equilibrium for (*±*)-blebbistatin concentration ranging from 0 to 160 *μ*M are given for comparison. The scale bar indicates 200 *μ*m. Variations of both surface tension (**d**) and Young’s modulus (**e**) with (*±*)-blebbistatin concentrations are reported. Inhibition curves are fitted for the surface tension (red curve) and the Young’s modulus (green curve) with a dose-response providing an *IC*_50_ value for the (*±*)-blebbistatin of 10 *±* 3 *μ*M and 7 *±* 5 *μ*M respectively corresponding to an *IC*_50_ for the (*−*) active enantiomer of blebbistatin around 6 *±* 2 *μ*M and 4 *±* 3 *μ*M respectively as the ratio of the active negative form is around 50-60%.

**Figure 3:**
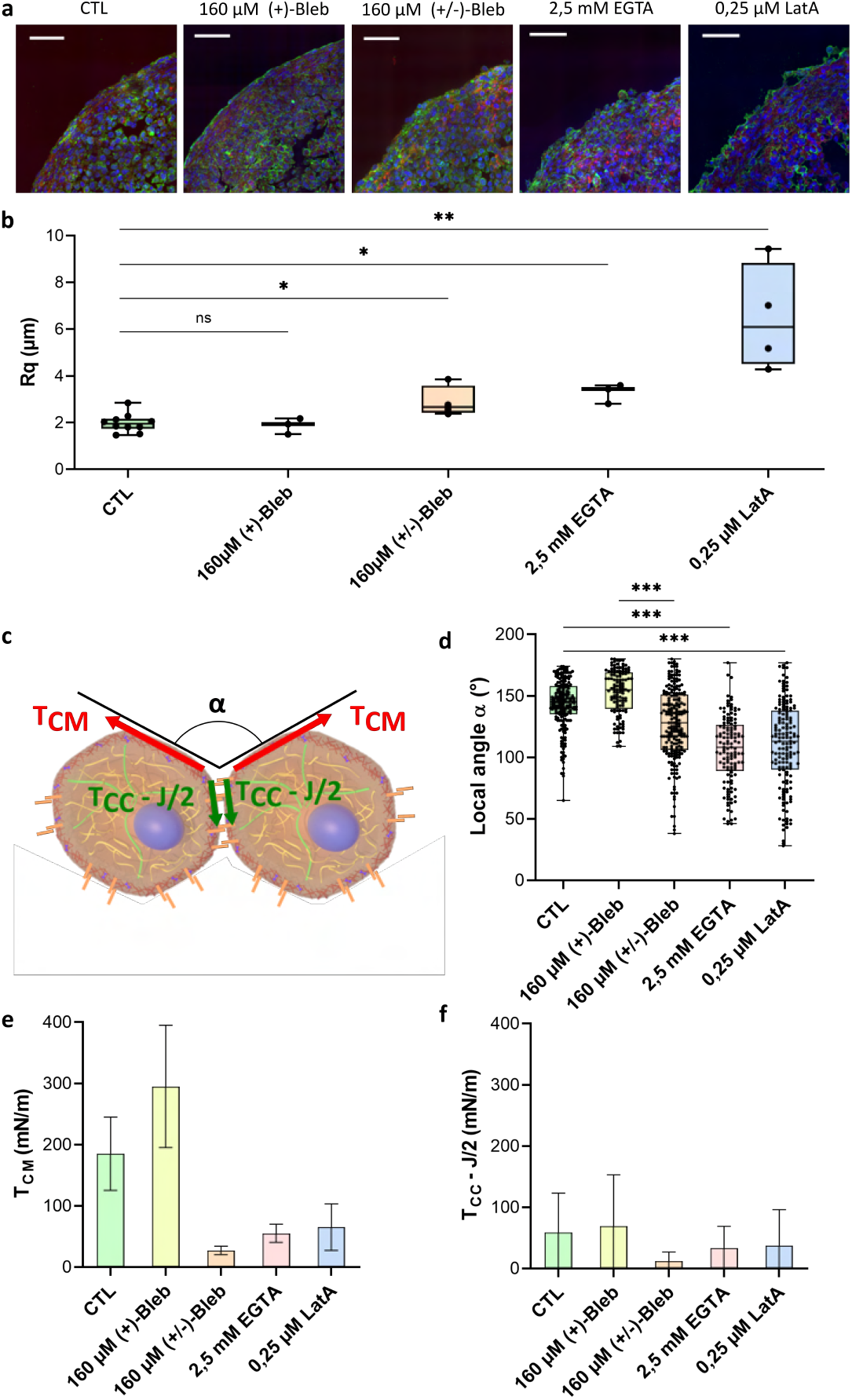
Geometrical analysis of cells at the aggregate surface. **a)** Immunofluorescence images of cryosections of multicellular aggregates obtained by magnetic molding with different conditions. Control cells are compared to aggregates produced with 160 *μ*M (+)-blebbistatin (inactive enantiomer), 160 *μ*M (*±*)-blebbistatin (active enantiomer), 2.5 mM EGTA or 0.25 *μ*M latrunculin A. DAPI is shown in blue, pan-cadherin in green, and F-actin in red. Scale bar = 50 *μ*m. **b)** Profile surface roughness parameter *R*_*q*_ (root-mean-squared) in each condition for at least *N* = 2 spheroids. **c)** Schematic of two neighbouring cells with a local contact angle *α* and the respective tension at the cell-medium interface *T*_*CM*_ and effective tension at the cell-cell contact *T*_*CC*_ *− J/*2. **d)** Contact angle between cell surfaces measured in each condition for at least *N* = 2 spheroids. **d-e)** Deduced values of the cell tension at the cell-medium interface (**d**) or of the effective adhesive tension at the cell-cell contact (**e**) in each condition.

**Figure 4:**
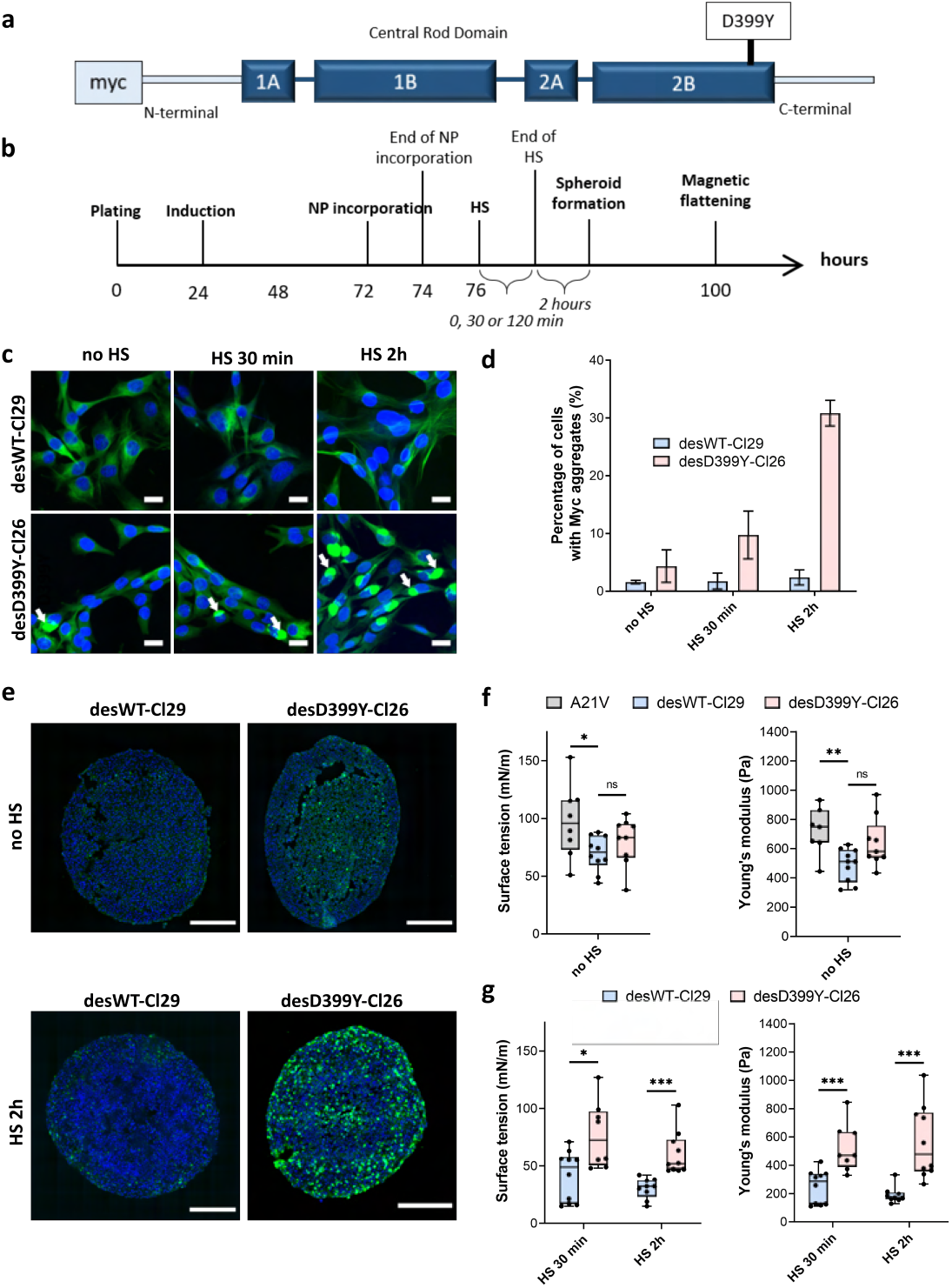
Protein aggregation of desmin (mutation D399Y) increases surface tension and Young’s modulus of C2C12 spheroids. **a)** Representation of desmin with the missense mutation D399Y located in Rod Domain. The expressed exogenous mutated desmin is Myctagged at the N-terminus (adapted from [41]).**b)** Experimental procedure. Desmin expression was induced 24 h after cell plating with doxycyclin for 48 h. Magnetically labelled cells (see Methods) experienced a heat shock (HS) for 0, 30 or 120 min before spheroids were molded. Spheroid surface tension and Young’s modulus were measured 3 days after expression induction. **c)** Immunofluorescence images of desWT-Cl29 (cells stably expressing exogenous desmin WT) and desD399Y-Cl26 cells (cells stably expressing exogenous mutated desmin) in 2D after 0, 30 or 120 min HS. DAPI is shown in blue and the tag Myc in green. Scale bar = 20 *μ*m. **d)** Percentage of cells with desmin protein aggregates for each condition. Desmin protein aggregation increases for desD399Y-Cl26 cells with the duration of the HS while it remains stable around 2% for desWT-Cl29 cells. Mean values represented with respective standard deviations and at least 3 independent experiments for each condition. **e)** Immunofluorescence images of multicellular aggregate cryosections of desWT-Cl29 or desD399Y-Cl26 cells with or without HS. DAPI is visible in blue, Myc-tag in green. The results are reminiscent of the ones in 2D. DesD399Y-Cl26 spheroids exhibit sparse aggregation without HS, enhanced by 2 h HS. Scale bar = 200 *μ*m. See Supplementary Figure 8 for zoomed images. **f)** Surface tension and Young’s modulus of A21V cells (control cells stably transfected with empty vector) spheroids compared with desWT-Cl29 and desD399Y-Cl26 cells to test for the influence of desmin overexpression. **g)** Surface tension and Young’s modulus of desWT-Cl29 and desD399Y-Cl26 spheroids with a HS of 0 (no HS), 30 or 120 min. (f-g) At least 3 independent experiments for each condition and *N* = 7 spheroids. Floating bars represent min to max variations and the midline indicates the median.

### Relation between macroscopic properties and molecular or cellular characteristics: a multi-contribution pattern

The molecular origins of spheroid surface tension are numerous. While differential adhesion hypothesis has been first evidenced by Steinberg [34] and related with the level of cadherins [30], the role of actin cortex was pointed out in individual cells [35], in spheroids [9] or embryogenesis [36] evidencing a differential interfacial tension hypothesis combining the influence of adhesion and cell surface tension [37]. By comparing energy of cells in the core of the aggregate to the one at the interface, the surface tension is given by:

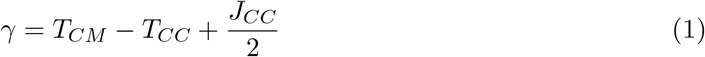

where *T*_*CM*_, *T*_*CC*_ and *J*_*CC*_ stand for the cortical tension at the cell-medium (CM) or at the cell-cell (CC) interface and the intercellular surface adhesion energy (*J*_*CC*_ *>* 0) respectively [9]. We check for this multi-parameter influence by looking at cell-cell contacts inhibition and at changes in actin structure or acto-myosin contractility. Intercellular adhesion is mediated by multiple cell-cell adhesion proteins among which cadherins play a key role. EGTA as a calcium chelator, reduces the efficiency of this homophilic adhesion and modulates cell-cell adhesion strength. Its addition leads to a more than 5-fold decrease of both Young’s modulus and surface tension (Figure 2c and d). The relationship between adhesion bond energy and the surface tension [30] is thus tested. An apparent proportionality between surface tension and Young’s modulus is obtained (Figure 5), reminiscent of the one observed between surface tension and elasticity of the whole aggregate (itself related to cortical tension) through co-regulation mechanisms [38]. Latrunculin A disrupts the actin filaments by binding to actin monomers thus precluding its polymerization [39], its addition gives some non-connected patches of actin filaments with a lack of long-range organization (Supplementary Figure 6). Young’s modulus as well as surface tension are dramatically decreased by its presence. Besides, its action on macroscopic properties are dose-dependent as seen by the comparison between 0.15 *μ*M (2-fold decrease) and 0.25 *μ*M (5 *−* 7-fold decrease) concentrations. Our results are consistent with the dependence previously noticed of surface tension and viscosity for latrunculin-treated cells [11]. They extend this dependance to stiffness.

**Figure 5:**
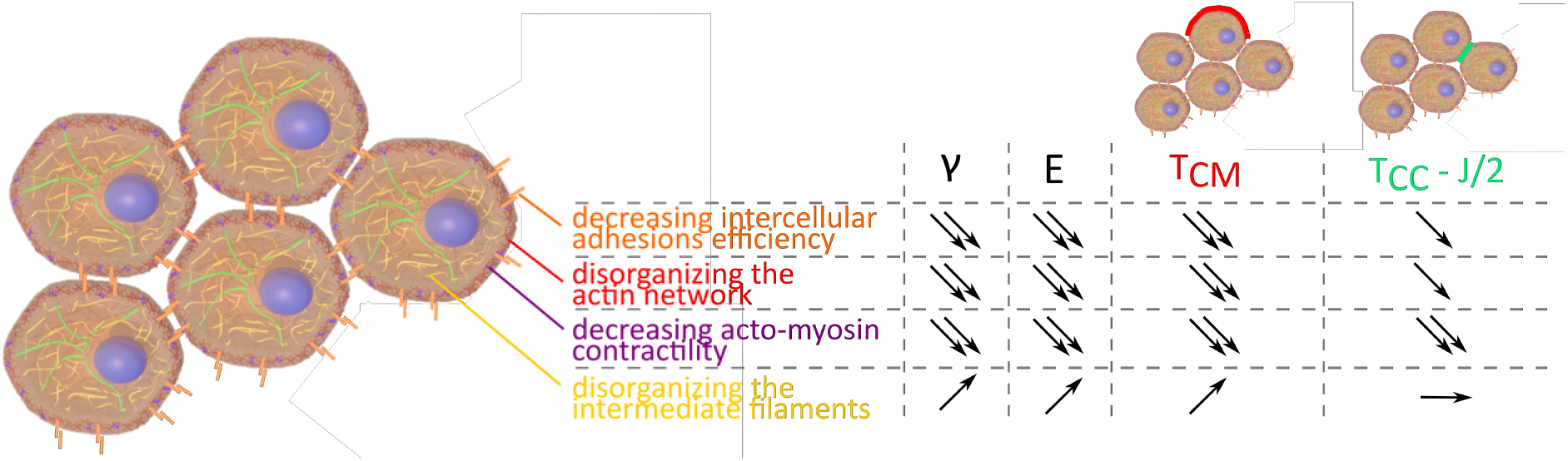
Evolution of *γ*, E, *T*_*CM*_ and *T*_*CC*_ *− J/*2 depending on intercellular adhesions, the actin network, acto-myosin contractility and intermediate filaments.

To further test the sensitivity of the magnetic tensiometer, we selected reagents with a wider accessible range still allowing cohesive spheroid formation (Supplementary Figure 7). (*−*)-Blebbistatin inhibits contractility by blocking myosins. Its inhibition potency is quantified by an *IC*_50_ in the range from 2 to 7 *μ*M depending on myosin type [40]. In our experiments, blebbistatin can be used on a wide range of concentrations (0 *−* 160 *μ*M). This inhibitor leads to increasing aggregate deformation with concentration (Figure 2c). Surface tension and Young’s modulus dramatically decrease with (*−*)-blebbistatin concentration (Figure 2d and e) with a 100-fold noticed decay. The extracted *IC*_50_ is around 6 *±* 2 *μ*M for surface tension and 4 *±* 3 *μ*M for elasticity reproducing the one obtained at the molecular level as the ratio of the active negative enantiomer is around 50 *−* 60%.

### Correlation with geometrical analysis: tensions at the interfaces

We were able to test the relation between surface cell shape and arrangement, and surface tension by looking at cell surface morphology on cellular aggregates cryosections (Figure 3a) upon the use of the different drugs. Latrunculin A and EGTA treated aggregates show rounded cells at the interface while control cells flatten on the surface without extending over multiple cells. (*±*)-Blebbistatin treated aggregates have an intermediate behavior. To quantify these observations, we extracted both the rugosity of the profile and the mean contact angle *α* between cells at the interface. They show similar variations. For control aggregates and (+)-blebbistatin aggregates, cells at the surface spread out, having a flat angle at the cell-cell contacts and a small rugosity length. Drug drastic effects can be noticed on EGTA and latrunculin A treated aggregates: rugosity increases (by 3 for the latrunculin A cells and by 2 for EGTA cells) while contact angles deviate from flat angle to get close to 100°. Overall, the surface tension variations are correlated with a change of morphology of the cells at the interface. As already noticed [37], high surface tension usually appears as the hallmark of flattened cells and low roughness. However, the case of (*±*)-blebbistatin inhibitor shows that multiple parameters have to be considered as the low surface tension obtained with the myosin inhibitor does not lead to rounded cells at the interface. Surface tension arises from a balance between cortical tension at the cell-medium and adhesion and cortical tension at the cell-cell interface (Equation 1) [37, 9]. Surface tension quantifies this interplay and not only the tension at the cell-medium interface while being predominant. All the considered aggregates do not show up elongated surface cells extending over multiple inner cells meaning that the effective adhesion 2*T*_*CC*_ *− J* is smaller than the double of the cortical tension at the cell medium interface. In this configuration, differential interfacial tension hypothesis and more sophisticated models provide similar results [37] and the local mechanical equilibrium at the three phases cell/cell/medium contact line gives the relation [9]:

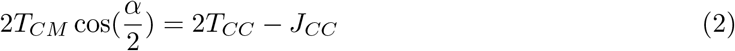

Both effective adhesion per cell *T*_*CC*_ *− J*_*CC*_*/*2 and cortical tension at the cell medium interface *T*_*CM*_ can then be deduced from surface tension and contact angle measurements. First we checked that the tension at the cell-medium interface is predominant in the value of surface tension. As noticed on the cryosections, EGTA and latrunculin A impact predominantly the tension at the cell-medium interface with a reduction by a factor around 3. Latrunculin A by depolymerizing actin reduces global cortical tension as well as the cell-cell adhesion by weakening cadherin anchorage through actin. Feedbacks between adhesion molecules and the cytoskeleton are indeed abundant and explain the lower variations of effective adhesion upon latrunculin A and EGTA addition due to a possible compensation of the tension decrease by an increase of adhesion.

Besides blebbistatin has an important effect on both effective adhesion and tension at the cell medium as it impacts neither intercellular adhesions nor the cytoskeleton structure but the ability of the cell cortex to contract. This inhibitor by decreasing tension at the cortex reduces both *T*_*CM*_ and *T*_*CC*_ thus affecting tension and effective cell-cell adhesion in a more drastic way than latrunculin or EGTA.

Hence the magnetic tensiometer appears as a highly sensitive tool to look at tissue model mechanics and its relation with modifications at the cellular level.

### Intermediate filaments action on macroscopic mechanical properties of muscular tissue models

Desmin is one of the main intermediate filaments in muscle cells where it plays an essential role in maintaining mechanical integrity and elasticity [42] at the single cell level, it stands as a marker for muscular cell differentiation but its role on tissue surface tension maintenance and elasticity has not been explored. Desmin mutations are involved in human diseases such as certain skeletal and cardiac myopathies [43], characterised histologically by intracellular protein aggregates containing desmin. We focus on the missense mutation D399Y [41]. In this cellular model, desmin aggregation is induced by heat shock in 2D culture (Figure 4a-d) but also in multicellular spheroids (Figure 4e). In desmin mutated cells, heat shock duration monitors the percentage of cells presenting desmin-aggregates with up to 30% for mutated desmin cells (desD399Y-Cl26) for 2 hours stress compared to a 2% ratio in the case of wild-type desmin expression (desWT-Cl29). Both surface tension and Young’s modulus are slightly decreased by the overexpression of desmin (wild-type or mutated) (Figure 4f). Desmin overexpression and protein aggregates (Figure 4e) may cause this decrease as desWT-Cl29 and desD399Y-Cl26 cells behave the same way. Focusing on exogenous desmin expressing cells, an increase in surface tension and elasticity dependent on the heat shock duration is observed in mutated desmin compared to wild-type desmin cells (Figure 4g): a 2-fold (1.8-fold) increase is noticed between desWT-Cl29 and desD399Y-Cl26 spheroid surface tension and a 3-fold (2-fold) increase for the stiffness for 2 hours (30 min) heat shock. Effects on mechanical properties are thus correlated with the percentage of cells containing desmin aggregates. Local cell disorganization modifies surface tension probably through individual cell tension modification.

Looking at the local geometry at the interface, cell-cell contact angle is slightly modified by the overexpression of mutated desmin and its aggregation (Figure 9a). Considering effective adhesion *T*_*CC*_ and tension at the cell-medium interface *T*_*CM*_, empty vector transfected cells (A21V) aggregates have the same behavior as control cell aggregates. Desmin mutants also show as control cells a predominance of tension at the cell medium interface (Supplementary Figure 9). Besides overall desmin overexpression (both wild-type and mutated) and protein aggregates influence the cortical tension at the cell-medium interface with an increase by a factor almost 2 between desWT-Cl29 and desD399Y-Cl26 spheroids. Conversely desmin aggregation on effective adhesion is smaller (a factor 1.4 is noticed).

## Discussion

Magnetic tensiometry achieves high sensitivity and precision even on a cell system that do not easily deform as muscle cells. Its sensitivity is in the range of the one obtained with compression plates tensiometer experiments for lower surface tension cell aggregates [44] while the accessible range of measurable surface tension is larger. The magnetic tensiometer measurements allows to provide dose-response curves thus representing an accurate tool to quantitatively characterize inhibitors action on mechanical properties. Its robustness is related to the spheroid formation technique that allows to provide well-controlled aggregates of unprecedented size with radius and content perfectly monitored in less than 24 hours. It offers a unique opportunity to explore mechanical properties even in a dose-dependant manner and to quantitatively extract the effects of mutations.

Measuring surface tension and elasticity proves to be a powerful tool to explore tissues mechanical properties. Both are key elements on tissue shape maintenance [45, 46]. Looking at 3D unorganized tissue models, we recapitulate the major cellular factors that influence tissue shape and response to perturbations. Intercellular adhesions and actin cortex structure or contractility appear unsurprisingly as fundamental as was found *in vivo* [47, 48]. Their decrease reduces both surface tension and elasticity (Figure 5). The major contribution of cell contractility in tissue shape maintenance is also pointed out regarding the resulting decay obtained for surface tension and stiffness at high concentration of blebbistatin. Inhibition at the microscopic level thus correlates with surface tension and elasticity evolution, confirming them as relevant indicators of molecular state and cellular contractility.

Altogether, our results suggest that measuring mechanical properties at tissue-scale provides insights into the molecular level [49] and that conversely, molecular modifications can induce mechanical changes at tissue-scale. Complemented by local measurements of the contact angle, our approach is able to give the relative influence of cytoskeleton structure, adhesion molecules and cortical tension on the surface tension and may distinguish between tension and effective adhesion modifications.

Desmin is the most representative intermediate filament for muscle cells. It has a pivotal role in myofibril architecture in mature muscle but also exerts important function on the adaptation of muscle cells to passive stretch and contractility [50]. Desmin organization defects impact severely muscle formation and maintenance and cause myopathies or cardomyopathies [23]. In single cells, desmin aggregation has been associated to sarcomere misalignment but also impact biomechanical properties. We measure its impact on surface tension and elasticity: desmin aggregation leads to an increase on both elasticity and surface tension. Desmin acts on muscle functioning and integrity [51, 52], its disorganization may impair individual cell contractility [53]. Our results shed light on its major role at the tissue level where it impacts cell tension mostly at the cell-medium interface. Intermediate filaments organization therefore appears as essential in muscle tissue shape maintenance and mechanical properties even at early stage of differentiation. At the individual scale, mutated desmin cells show modified reorientation dynamics [54] and are stiffer [42], a behavior reminiscent of the elasticity increase noticed on mutated desmin filaments D399Y [55]. This stiffening is slightly translated at the multicellular scale on the Young’s modulus (Figure 4f). Moreover while desmin overexpression slightly decrease surface tension (Figure 4f), desmin protein aggregation increases its value. Protein aggregates by more rigid network nodes strenghten cell cortex [42] and may reinforce effective cortex tension thus increasing tissue surface tension.

Surprisingly, regarding desmin aggregation, no effect on individual cell stiffness was reported but rather a distribution modification [42]. Conversely we demonstrate in muscle precursor cell spheroids that desmin aggregation impacts both elasticity and surface tension pointing out the importance of desmin organization on macroscopic tissue mechanics even at early stage. Surface tension and elasticity appear thus as good reporters of the individual cellular state but also of the multicellular organization. Two reasons may be involved. First around 150 000 cells constitute spheroids. Measuring macroscopic properties thus provides an averaging over a large population of the individual cell properties; secondly collective enhancements may arise as modifying cell organization modulates cell-medium and cell-cell tensions both implicated in surface tension [9].

## Conclusion

Macroscopic properties of multicellular spheroids such as surface tension and elasticity appear as highly sensitive markers for cell cortex and cell-cell adhesion modifications. Measuring them by an approach based on magnetic cell labelling and multicellular aggregates tensiometry allows to explore dose-response evolution correlated with microscopic inhibition potency. The precision provided by the magnetic tensiometer opens a new field of investigation to test the impact of potential drugs or genetic modifications on the mechanics of early-stage tissue models. This approach was tested on a cellular model of desmin-related myopathies and demonstrated that desmin disorganization induces macroscopic changes in early-stage tissue models, undetected from now at the individual cellular scale. The fully integrated system of magnetic molding and magnetic tensiometry can be envisioned as a powerful tool for the study of fundamental biological processes and for the detection of mechanical effects leading to a better understanding of skeletal muscle dystrophies.

## Methods

### C2C12 cell culture

C2C12 WT (ATCC CRL-1772) were cultured in Dulbecco’s modified Eagle’s medium (DMEM, Gibco), supplemented with 1% Penicillin-Streptomycin (P/S, Gibco) and 10% Fetal Bovine Serum (FBS, Gibco). C2C12 A21V (stably transfected with empty vector), desWT-Cl29 (stably expressing exogenous human WT desmin) and desD399Y-Cl26 (stably expressing exogenous human D399Y desmin) were cultured in 1% P/S, 20% FBS, 1 mg/mL Geneticin (10131027, Gibco) and 2 *μ*g/mL Puromycin (P7255, Sigma-Aldrich) as described in Segard et al. [41].

### Magnetic labelling

Iron oxide superparamagnetic nanoparticles (8 nm diameter) (provided by PHENIX laboratory, UMR 8234, Paris), were obtained by alkaline coprecipitation followed by an oxidation into maghemite according to Massart procedure [56]. Finally, the aqueous solution was stabilized electrostatically by the adsorption of citrate anions on the nanoparticles surface.

Cells were magnetically labelled thanks to an incubation of 2h with a solution of iron oxide nanoparticles at [*Fe*] = 4 mM and supplemented with 5 mM citrate in RPMI medium (Gibco). The viability and the proliferation of cells after magnetic labelling were assessed by Alamar Blue showing no impact of the magnetic labelling right after the labelling or after one day (Supplementary Figure 1).

### Magnetic molding

Labelled C2C12 cells were incubated for at least 2h in complete medium. Agarose molds were previously prepared by heating a solution of 2% agarose (A0576, Sigma-Aldrich) in phosphate buffer saline (PBS) up to boiling. The resulting solution was then poured into a 60 mm culture Petri dish containing 1.6 mm steal beads (BI 00151, CIMAP) held in place by magnets below the Petri dish. Once the agarose solidified, the beads were removed carefully and the agarose wells were coated with a non-adhesive coating solution (07010, Stemcell Technologies) for 30 min at room temperature (RT). Cells were then detached with 0,05% Trypsin-EDTA and seeded in the wells thanks to magnet attraction (1.5 10^5^ cells per well). Spheroids were then incubated overnight at 37^*circ*^C, 5% *CO*_2_ in complete medium in normal conditions. At this point, the culture medium may be modified to add inhibitors (Figure 1). Spheroids were then extracted from the wells by gently pipetting the surrounding liquid. Spheroids of 540 *±* 50 *μ*m radius (N = 142) were obtained.

By measuring the magnetic moment of the aggregate with Vibrating-Sample Magnetometer (VSM) measurements (PPMS - Quantum Design) the magnetic moment per unit volume of the aggregate *M*_*v*_ can be determined and the value of surface tension deduced. *M*_*v*_ measurements ranged from 150 to 500 *A*.*m*^*−*1^ and were determined for each experiment. The force per unit of volume exerted on the spheroids ranged from 2.5 10^4^ to 8.5 10^4^ N.m^*−*3^.

### Magnetic force tensiometer

The magnetic force tensiometer is composed of a temperature-regulated tank at 37^*circ*^C sealed at the bottom and the sides by glass slides, a 6×6 mm cylindrical neodymium permanent magnet (S-06-06-N, Supermagnete, *B* = 530 mT, *grad*(*B*) = 170 T/m, averaged value between 100 *μ*m and 1.5 mm from the surface of the magnet), a lifting stage to approach the magnet and a camera (QICAM FAST1394, QImaging) equipped with a 1.5x zoom lens (MVL6X12Z, Thorlabs). The temperature regulated tank is filled with transparent culture medium (DMEM, high glucose, HEPES, no phenol red, Gibco). The bottom slide is treated with a non-adhesive solution (07010, Stemcell Technologies) to guarantee non-wetting conditions for the multicellular spheroids. Horizontality of the bottom slide has to be carefully checked. The aggregate side profile is registered with the camera. The magnet is approached at 100 *μ*m from the bottom of the aggregate. The macroscopic mechanical properties of the spheroid are obtained by the spheroid profile at equilibrium (10 min under magnetic flattening to ensure that equilibrium is reached). Surface tension is measured by fitting the spheroid profile at equilibrium with Laplace profile while Young’s modulus is determined thanks to the radius of the contact zone *L* and using Hertz theory as described in [28]. The Young’s modulus *E* equals 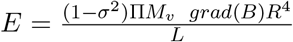 where *σ* stands for the Poisson ratio (*σ* = 1*/*2), *M*_*V*_ represents the magnetic moment per unit volume, *R* and *L* are the initial radius and the radius of the contact zone respectively. Briefly for the surface tension measurements, theoretical profiles were obtained by integrating numerically the Laplace law for capillarity and minimizing the quadratic error on the height *h*, the width *w* and the volume *V* of the spheroid (Supplementary Figure 3) to extract the capillary constant 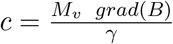 [14].

Depending on conditions, flattening occurs either with a two times regime or exhibit a single time decay. Spheroids have two ways of deformations to reach their capillary-driven equilibrium shape : first a rapid elastic deformation then a more viscous fluid behavior. This competition is driven by the size of the aggregate meaning that above a critical radius *R*_*C*_ [28], elastic deformation is complete and may be followed by viscouslike behavior to reach capillary-driven limit. We carefully checked that the radius of the spheroid is above this limit to be able to extract Young modulus. This modulus is extracted from the first step deformation while the surface tension is deduced from equilibrium shape. Tensio𝕏 is a dedicated MATLAB generated application freely available to extract both surface tension and Young’s modulus from initial and flattened profiles. Images of the initial and final state are first downloaded. User has to define the initial radius, the height and the width by pointing left/right and top/bottom frontiers of the spheroid on the two images. The radius of the contact zone is also extracted. The scale factor, the magnetic force per unit volume and an estimated surface tension value are entered. Obtained fit is then superimposed on the flattened image to check for its relevance and the deduced Young’s Modulus and surface tension are registered in a separated data file.

### Measurements of mechanical properties of C2C12 spheroids In drug conditions

For inhibitor conditions, the drug was added in the medium when the cells were seeded in the wells. *C2C12 spheroids were then incubated overnight at 37*^*circ*^C, 5% *CO*_2_ in DMEM 10% FBS, 1% P/S supplemented with the chosen concentration of EGTA (03777, Sigma-Aldrich), Latrunculin A (L5163, Sigma-Aldrich), (+)-Blebbistatin (203392, Sigma-Aldrich) or (+/-)-Blebbistatin (203390, Sigma-Aldrich). For (+/-)-Blebbistatin, inhibition curves were fitted with the function 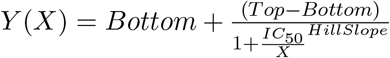 and with no constraints applied on the 4 variable parameters.

### C2C12 expressing mutated desmin

At day 0, cells were plated at 3000 cells*/*cm^2^ (2.25 10^5^ cells seeded in a T-75 flask). After 24h of incubation at 37°C, 5% *CO*_2_, expression of exogen desmin was induced by supplementing the culture medium with 10 *μ*g/mL doxycylin (D9891, Sigma-Aldrich) for 2 days. The medium was replaced with fresh medium every 24h. Iron oxide nanoparticles were incorporated into the cells at day 3. After 2 hours of recovery, a heat shock (HS) with a water bath at 42^*circ*^C for 0, 30 or 120 min was applied to the cells. 2 hours later, cells were then detached and used to form spheroids by magnetic molding or seeded in 24-well plates for desmin aggregation evaluation. After an overnight incubation at 37°C, 5% *CO*_2_, spheroids were magnetically flattened to measure their mechanical macroscopic properties, then fixed for 1 h at room temperature in 4% paraformaldehyde (PFA, J61899, Alfa Aesar). The corresponding cells grown in 2D in the 24-well plates were fixed for 15 *−* 20 min at RT in 4% PFA, to assess the percentage of cells with Myc aggregation for each condition.

### Cryosections and immunofluorescence

Spheroids were fixed in 4% PFA for 1 h at RT and stored in PBS at 4°C. For cryosections, spheroids were embedded in OCT (Optimal Cutting Compound, 361603E, VWR) for 1h at room temperature, they were then frozen in iso-pentane (24872.260, VWR), cooled down in liquid nitrogen, and then stored at *−*20°C. 6-10 *μ*m cryosections were obtained (Leica CM1520). For immunofluorescence labelling, cryosections or fixed cells were permeabilized 15 *−* 20 min in 0,1% Triton X-100 at RT. Non-specific interactions were prevented by an incubation with 5% BSA (#05479, Sigma-Aldrich) for 1 h at RT. Pan-cadherin (rabbit anti-pan cadherin (dilution 1:100 in PBS 0, 5% BSA, C3678, Sigma) for 2h at RT), and phospho-myosin (rabbit phospho-myosin light chain 2 (Thr18/Ser19) antibody (dilution 1:50 in PBS 0, 5% BSA, #3674, Cell Signaling) for 2 h at RT) were labelled. Both primary antibodies were coupled with an Alexa Fluor 488 goat anti-rabbit secondary antibody (dilution 1:500 in PBS 0,5% BSA, #4412, Cell Signaling Technology) for 2h at RT. Myc was labelled using mouse c-Myc (9E10) Alexa Fluor 647 (dilution 1:100 in PBS 0,5% BSA, sc-40AF647, Santa Cruz Biotechnologies) for 2 h at RT. F-actin was labelled using SiR-actin or SPY555-actin (dilution 1:1000 in PBS, Spirochrome) for 1h30 at RT, while nuclei were labelled with DAPI (dilution 1:1000 in PBS, D9564, Sigma-Aldrich) for 15-20 min at RT. All the samples were mounted with Fluoromont (F4680, Sigma-Aldrich) and stored at 4°C after gelation of the mounting medium at RT. Cryosections were imaged either on a Nikon microscope A1r25HD with a 100x oil objective or on an Axio observer Zeiss microscope equipped with a CSU-X1 Spinning disk with a 63x oil objective.

### Roughness measurements

The contour profile of the spheroids was extracted manually with Fiji (ImageJ) thanks to immunofluorescence images of spheroid cryosections (pan-cadherin, F-actin or phospho-myosin). The extracted experimental profile was then fitted by a circular arc and the roughness parameter *R*_*q*_ was measured by computing for each experimental point the distance *z* to the circular arc and then calculating 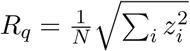 with *N* the number of points of the experimental contour. The smaller *R*_*q*_ is, the less rough the surface of the spheroid is, and the closer it is to a circular arc.

### Local contact angle measurements

Local contact angles were measured thanks to pan-cadherin immunofluorescence images of spheroid cryosections. Contact angles between cells at the spheroid surface were measured on Fiji (ImageJ) and pan-cadherin labelling was also used to confirm the adhesion between two neighbouring cells for each measurement. Measurements were performed on at least 2 spheroids for each condition and angles were measured all over the surface of the spheroid.

### Statistical analysis

All statistical tests were performed with a two-sided Mann-Whitney U test (Wilcoxon test) with MATLAB. p-value is used to indicate the statistical significance of the results (*, ** and *** correspond to *p <* 0.05, *p <* 0.01 and *p <* 0.001 respectively).

## Acknowledgements

The authors thank Alexandre Fromain for his help on VSM measurements and Ali Abou Hassan for providing us magnetic nanoparticles. This work was supported by the Program Emergence de la Ville de Paris (Grant MAGIC Project). The study was partially supported by the Labex Who Am I?, Labex ANR-11-LABX-0071, the Universitéde Paris, Idex ANR-18-IDEX-0001 funded by the French Government through its Investments for the Future program, by the AFM (french association for myopathies) AFM-22956, and by the French Defense Procurement Agency (DGA-AID) France. We acknowledge the ImagoSeine core facility of the Institute Jacques Monod (member of the France BioImaging, ANR-10-INBS-04).

## Data availability

Data supporting the findings of this study are available within the article and its Supplementary information files, and from M.R. upon reasonable request. Computing codes and resources should be found on M.R. webpage or requests should be addressed to M.R.

## Author contribution

I.N., F.D., S.H., S.B.P. and M.R. conceived the idea; I.N., F.D., and M.R. performed the experiments; I.N. and M.R. created the TensioX MatLab application; C.W, S.B.P. and M.R. supervised the study; I.N. and M.R. wrote the manuscript.

## Supplementary Figures

**Figure 1:**
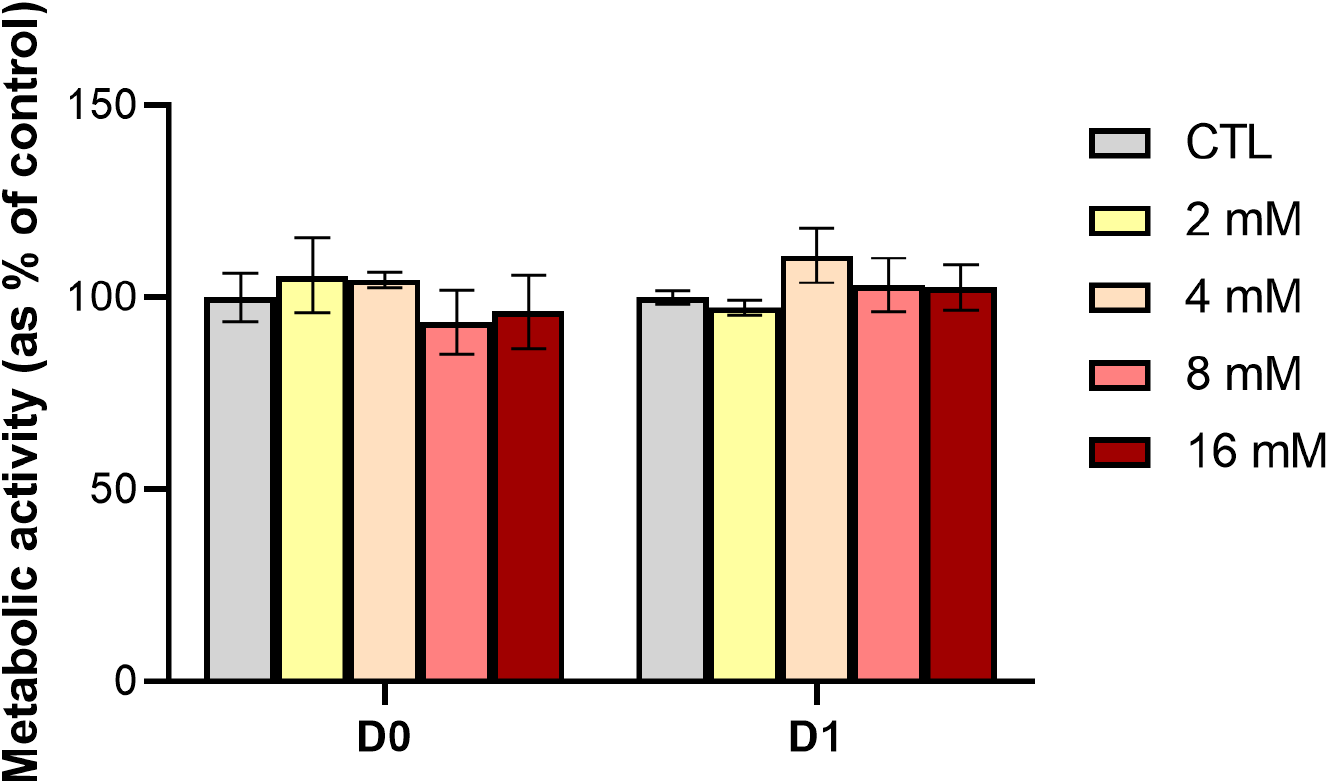
Metabolic activity for unlabelled (CTL) and labelled cells ([Fe] = 2 to 16 mM and 2 hours incubation time), 2 hours (D0) and one day (D1) after nanoparticle incorporation. To assess metabolic activity, the metabolic test Alamar Blue was used. Fluorescence was measured at *λ*_*exc*_ = 570 nm and *λ*_*em*_ = 585 nm. Values are interpreted relative to control values (unlabelled cells in complete medium, CTL) obtained under similar conditions. No influence of nanoparticle incorporation was observed on the metabolic activity and on the cell viability at D0 or D1 and for concentrations up to 16 mM.

**Figure 2:**
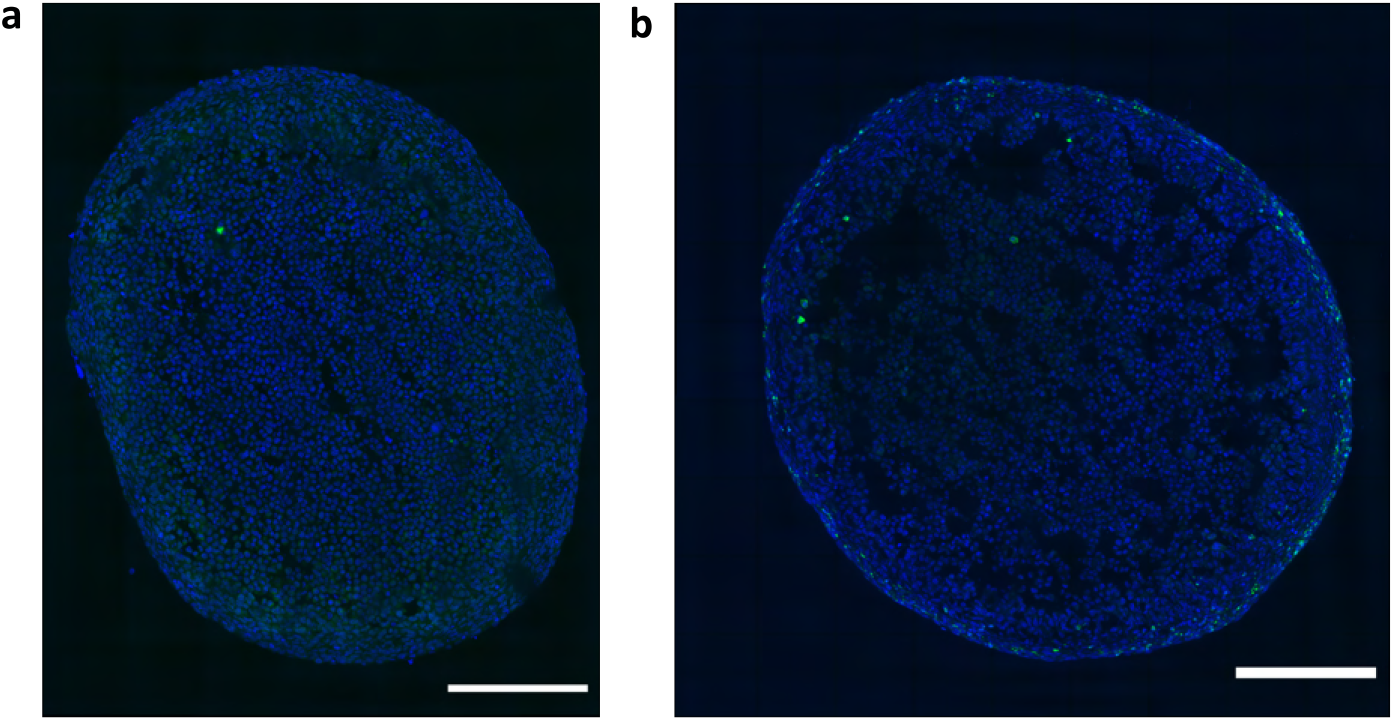
Immunofluorescence images of C2C12 spheroid cryosections to confirm non-hypoxic and non-apoptotic conditions in spheroids. **a)** DAPI is shown in blue and HIF*α* in green. HIF*α* is almost absent from the images. **b)** DAPI is shown in blue and cleaved caspase-3 in green. Only 2% of cells show a positive signal to cleaved caspase-3. Scale bars = 200 *μ*m.

**Figure 3:**
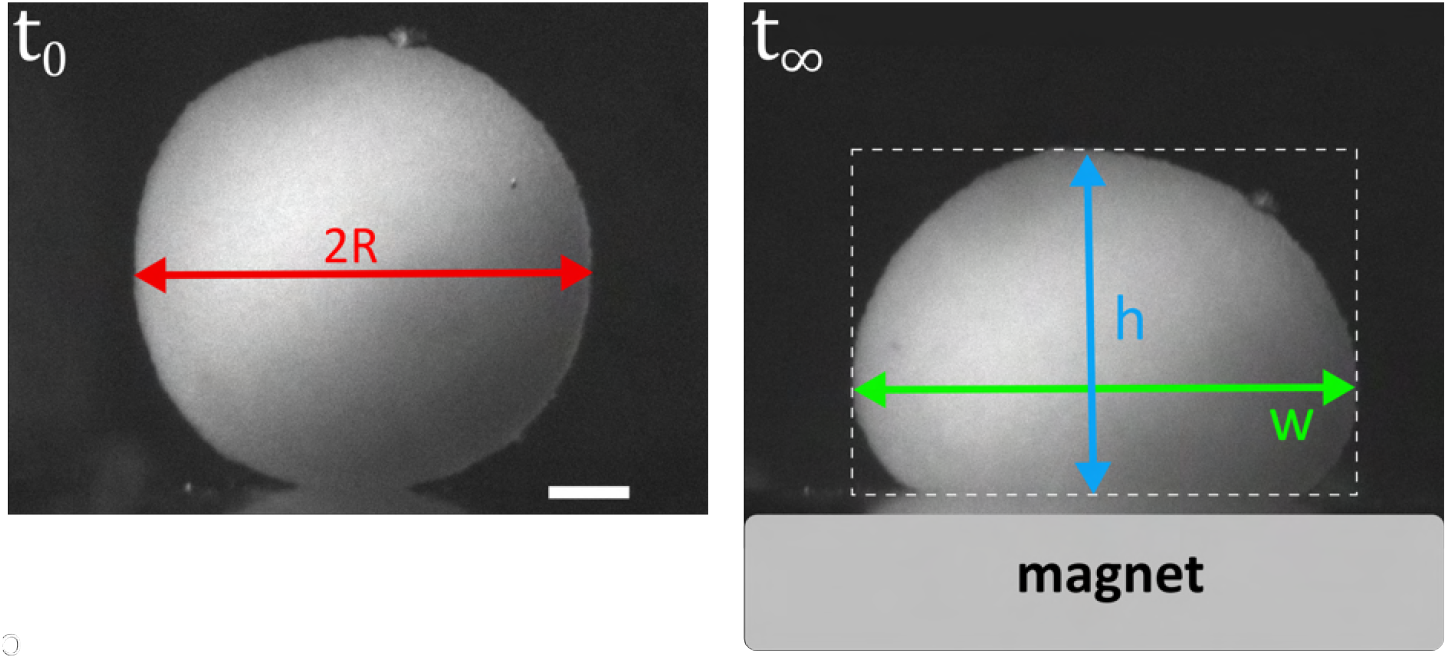
Extraction of experimental volume, width and height of the spheroid. The experimental volume V is calculated from the radius R of the spheroid at *t*_0_. The experimental width w (in green) and height h (in blue) are measured from the spheroid equilibrium profile. All these measurements are used in the minimisation program of the Tensio𝕏 application.

**Figure 4:**
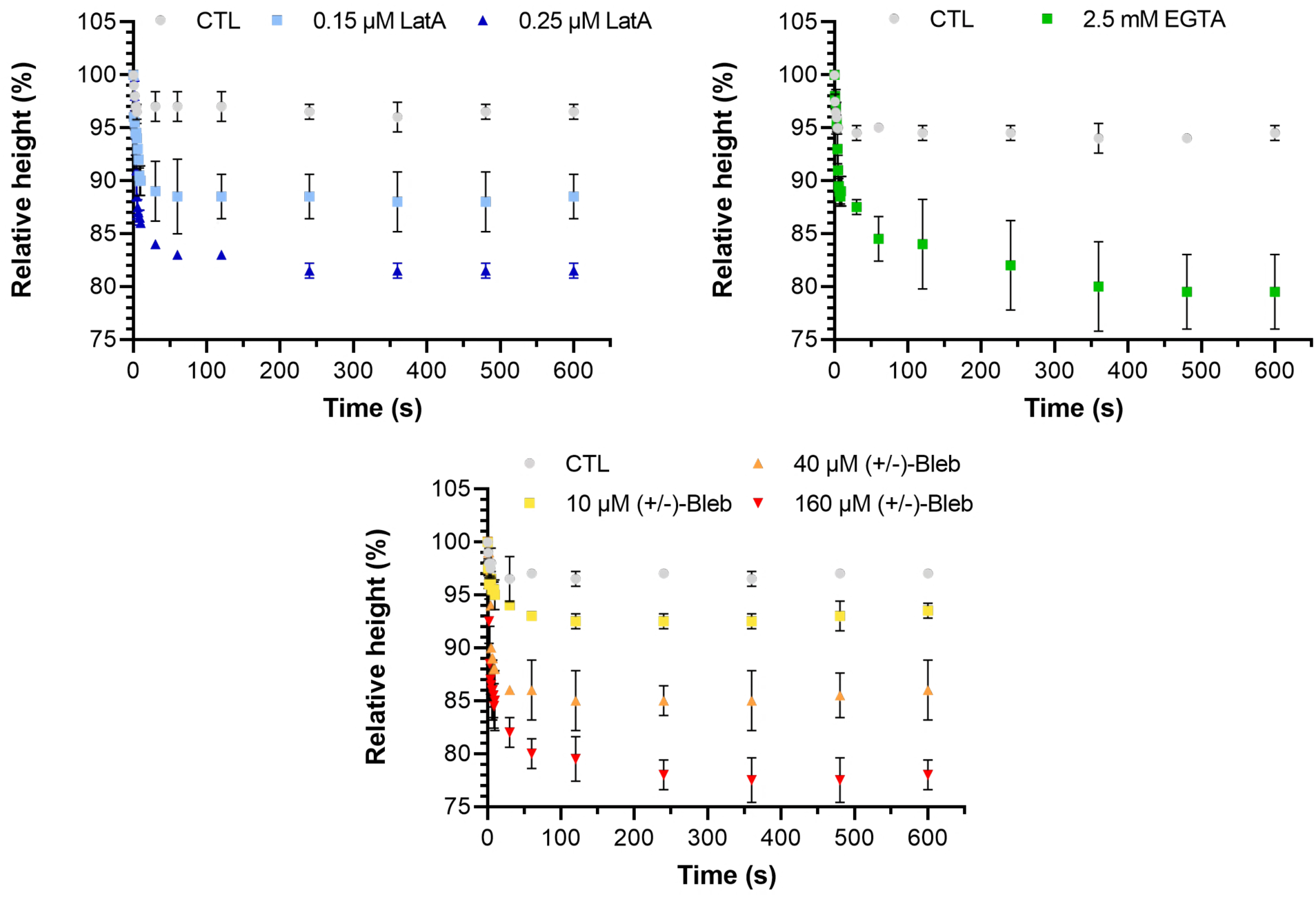
Relative height of the aggregate as a function of time for control, Latrunculin A (LatA), EGTA and (*±*)-Blebbistatin (*±*)-Bleb) conditions. At *t* = 0 s, the magnet is approached and the height of the spheroid is monitored over time. After 10 min of flattening, the equilibrium shape is reached and the height of the spheroid remains stable for each condition. Each curve corresponds to at least *N* = 2 spheroids. Mean values are represented with respective standard deviations.

**Figure 5:**
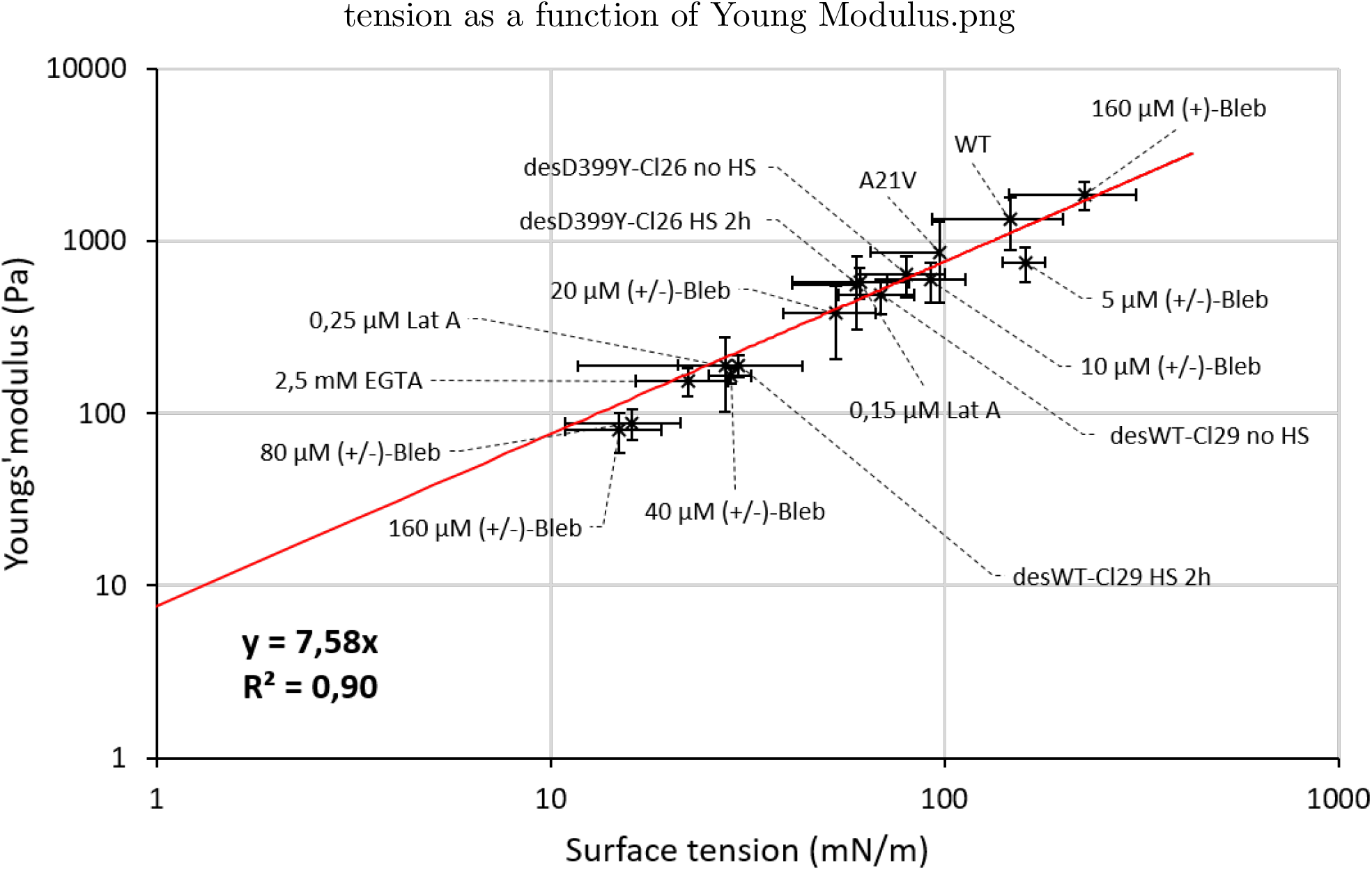
Young’s modulus of C2C12 spheroids as a function of their surface tension. Young’s modulus and surface tension are proportional across all the conditions tested for C2C12 spheroids. Values are represented in logarithmic scale.

**Figure 6:**
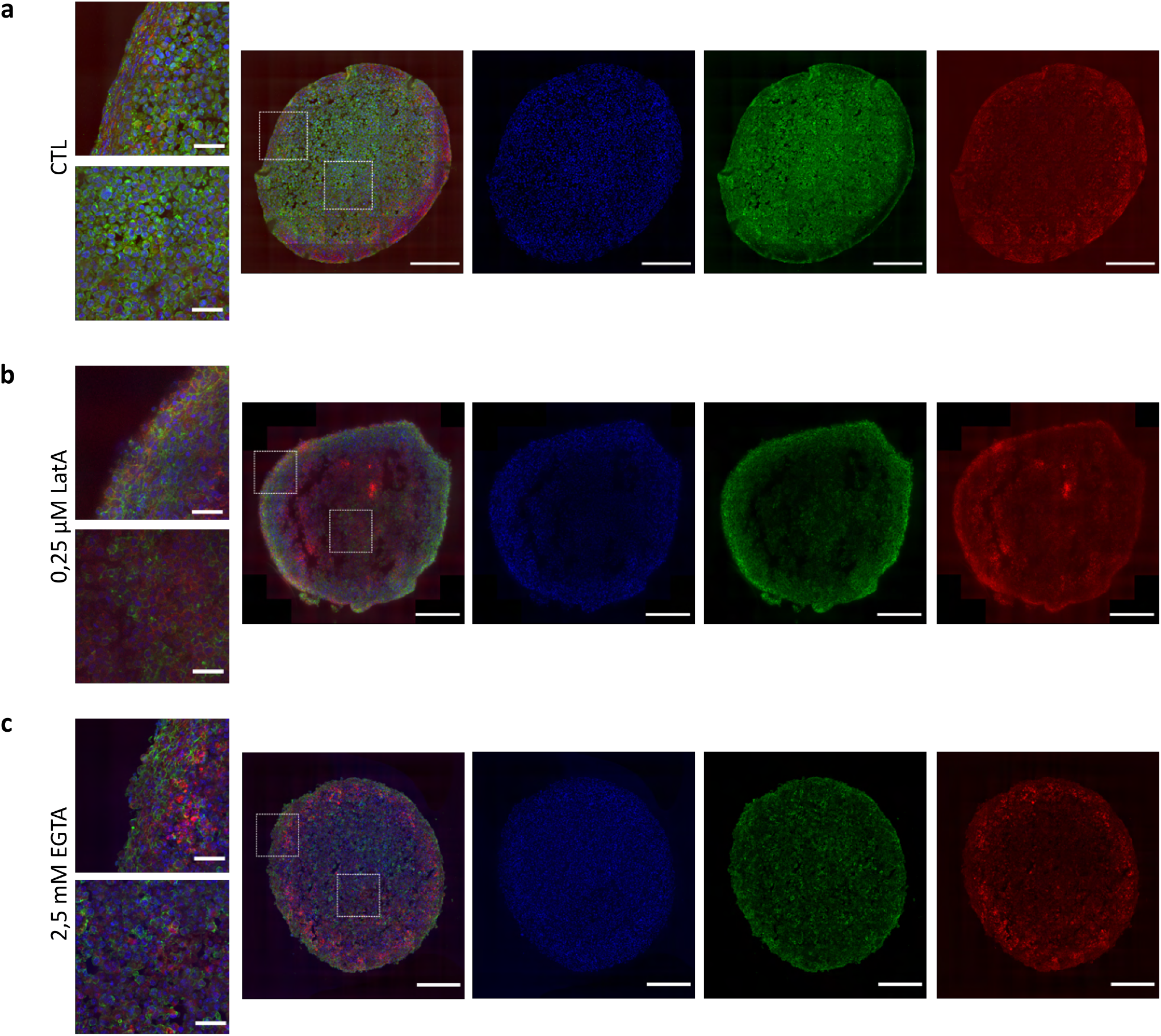
Immunofluorescence images of C2C12 spheroid cryosections for control conditions. **(a)**, 0.25 *μ***M Latrunculin A (b) and** 2, 5 **mM EGTA (c)**. DAPI is shown in blue, pan-cadherins are in green and F-actin is shown in red. Scale bar = 200 *μ*m and scale bar = 40 *μ*m for zoomed images.

**Figure 7:**
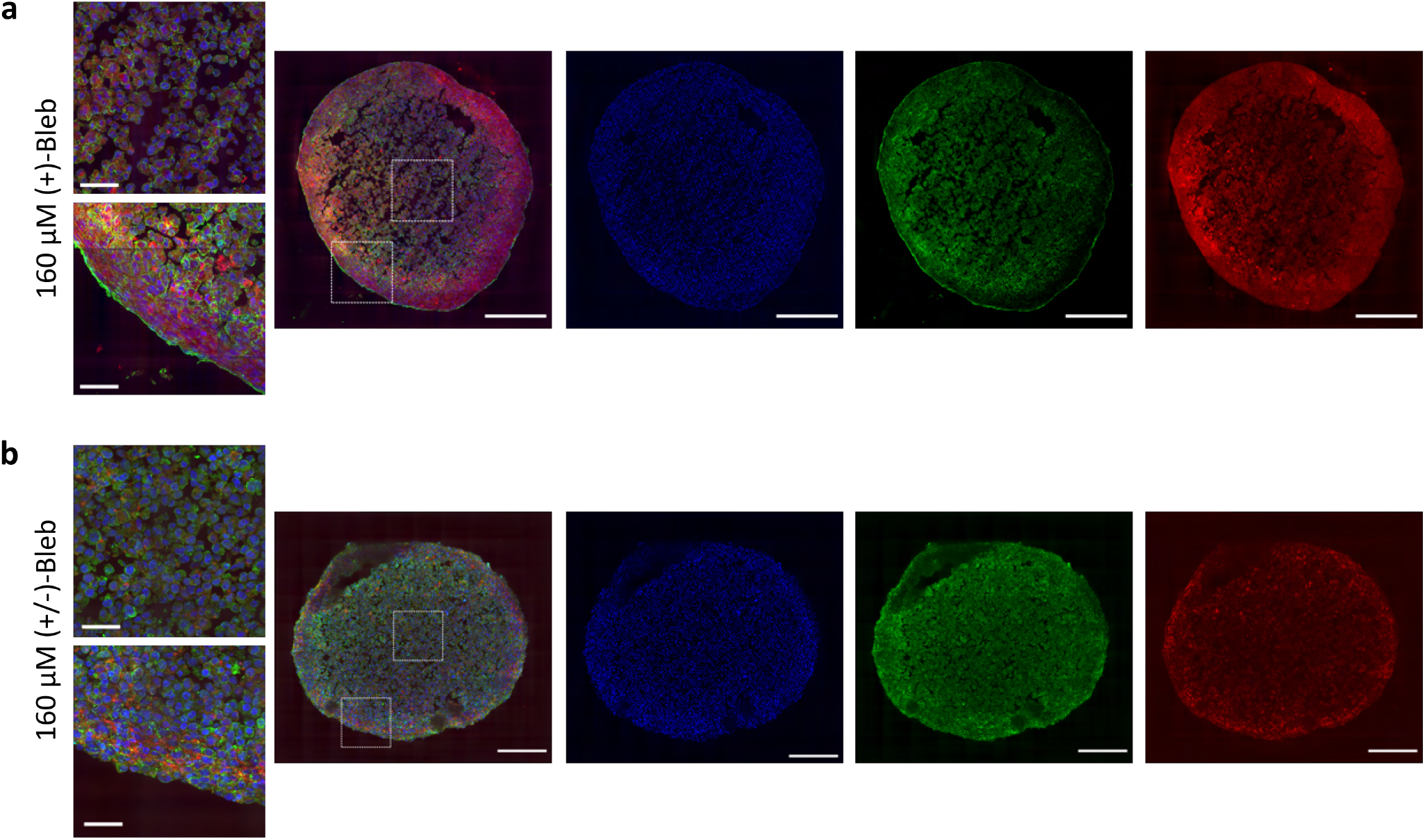
Immunofluorescence images of C2C12 spheroid cryosections for 160 *μ*M (+)-Blebbistatin (a) and 160 *μ*M (*±*)-Blebbistatin (b). DAPI is shown in blue, pan-cadherins are in green and F-actin is shown in red. Scale bar = 200 *μ*m and scale bar= 40 *μ*m for zoomed images.

**Figure 8:**
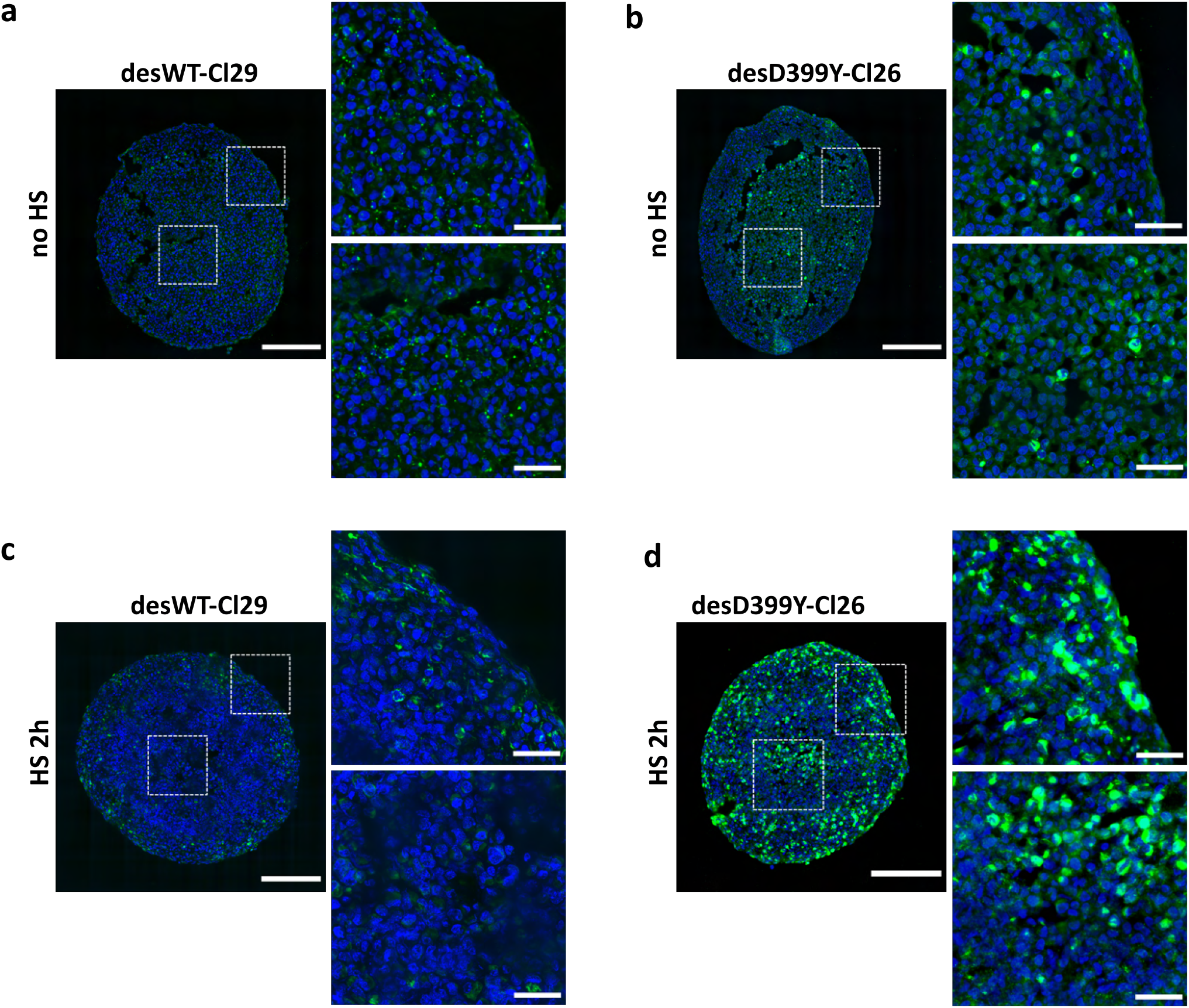
Immunofluorescence images of C2C12 spheroid cryosections of desWTCl29 (a,c) and desD399Y-Cl26 cells (b,d) for no HS (a,b) or 2 hours HS (c,d). DAPI is shown in blue and Myc is shown in green. Scale bar = 200 *μ*m and scale bar = 40 *μ*m for zoomed images.

**Figure 9:**
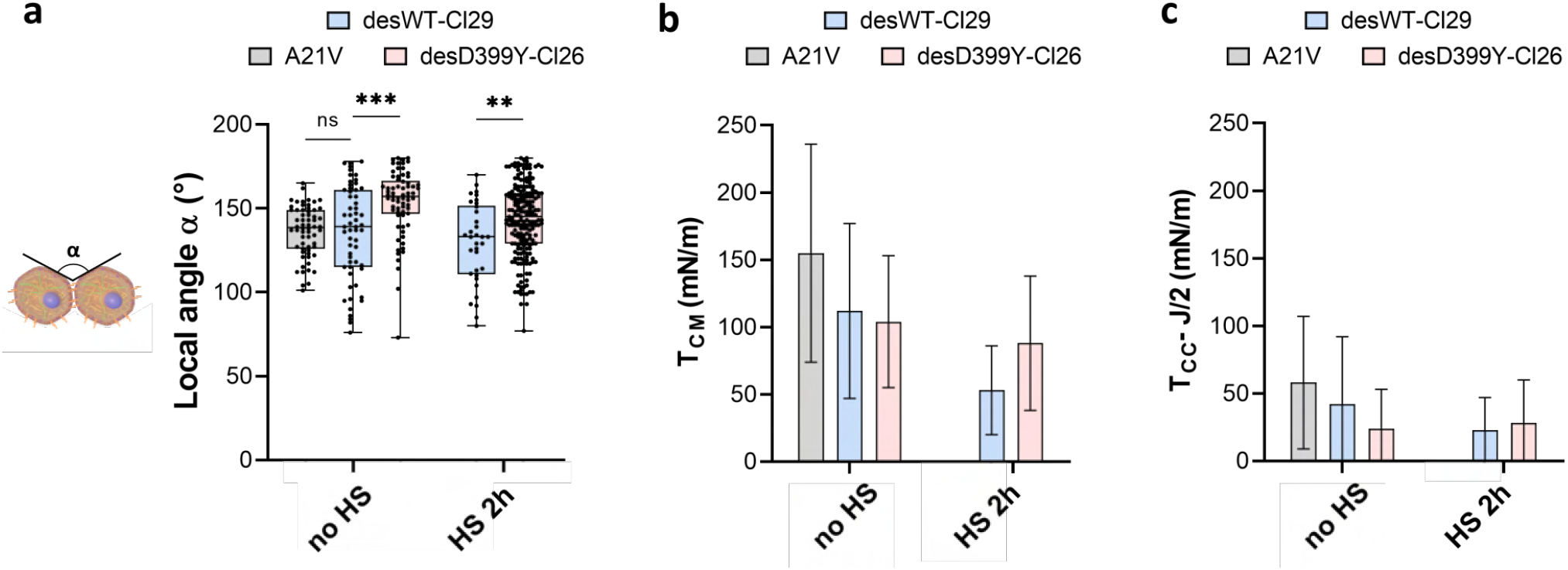
Geometrical analysis of cells at the aggregate surface for C2C12 A21V, desWT-Cl29 and desD399Y-Cl26 spheroids. a) Local contact angle between cells at the surface measured in each condition. **b-c)** Deduced values of the cell tension at the cell-medium interface (**b**) or of the effective tension at the cell-cell contact (**c**) in each condition.

## Application tutorial

### I. Installation

Tensio *X* can be installed and downloaded from **XXXXXX**. The application is coded in MATLABand the source code can be obtained at the same location.

No MATLAB license is required to run the application but MATLAB Runtime is required and is installed simultaneously with the application.

### II. Introduction and principle

This application is designed to measure the surface tension and the Young modulus of magnetic multicellular spheroids by flattening them with an external magnet as described in [1].

Briefly, iron oxide superparamagnetic nanoparticles (γ -Fe_2_O_3_) obtained via Massart’s procedure are incorporated into the cells then spheroids are formed by magnetic molding (overnight formation). Finally, spheroids are flattened with an external magnet and the side profile of each spheroid is monitored with a camera. The equilibrium shape of the spheroid is determined by the competition between surface tension and magnetic forces. Surface tension is measured by fitting the spheroid side profile at equilibrium while Young modulus is determined thanks to the radius of the contact zone using Hertz theory.

To obtain the measurements, four elements are needed:

- A picture of the spheroid at *t*_0_ (the initial spheroid needs to be spherical to ensure the accuracy of the measurements).
- A picture of the spheroid at *t*_f_ when the equilibrium shape of the spheroid under flattening is reached.
- The scale factor of the imaging system in m/pixels.
- The magnetic force per unit of volume applied on the spheroid *f*_v_ = *M*_v_ × *grad*(*B*), with *M*_v_ the magnetic moment per unit of volume (that can be measured using VSM, Vibrating Sample Magnetometry, measurements for example) and *grad*(*B*) the magnetic field gradient of the external magnet at the position of the spheroid.

### III. Step-by-step tutorial

- Open TensioX.
- Select the spheroid image at *t* = *t*_0_.

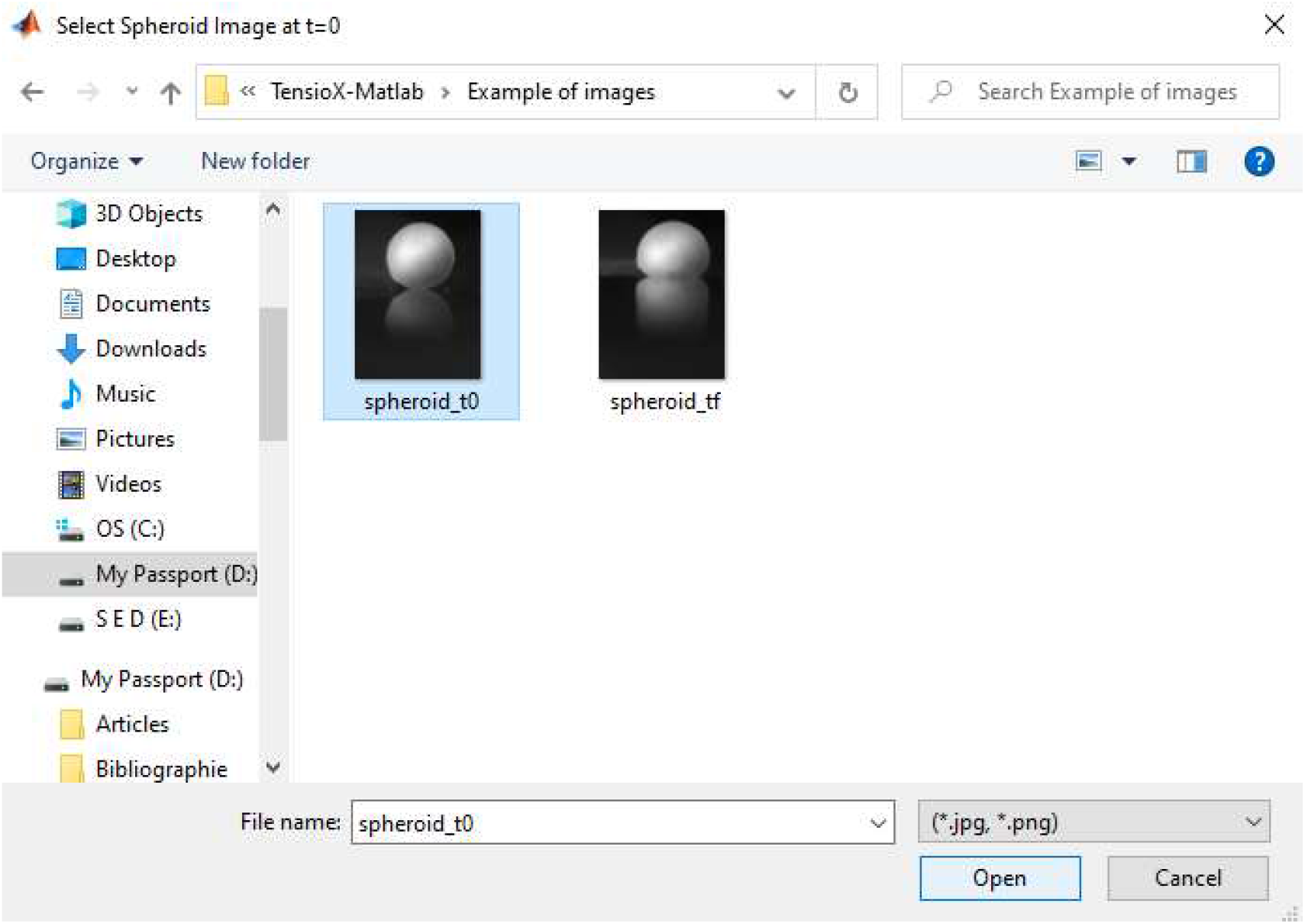
- Then select image of spheroid at *t* = *t*_*f*_ when the equilibrium shape is reached.

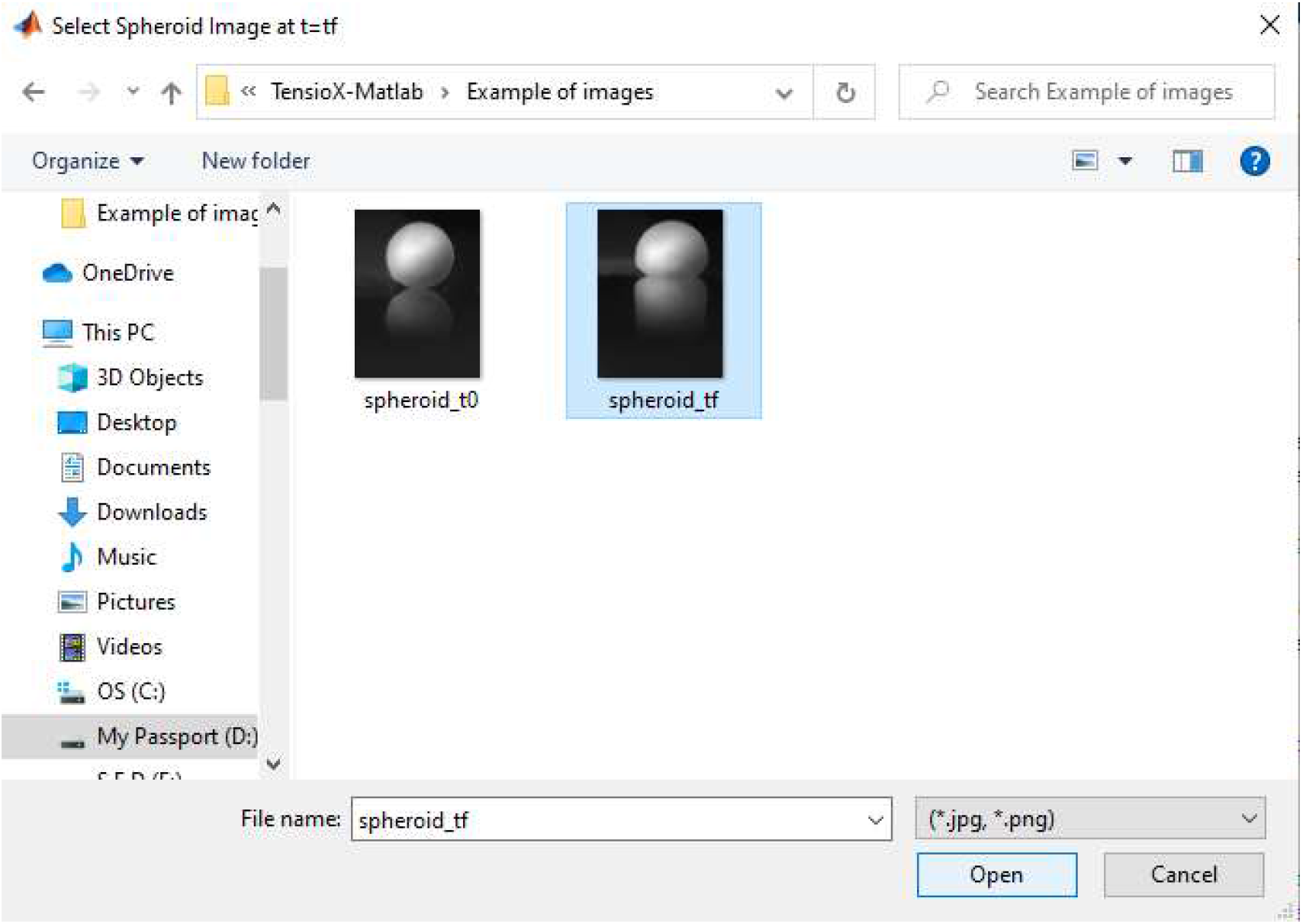
- Select folder where results will be saved.

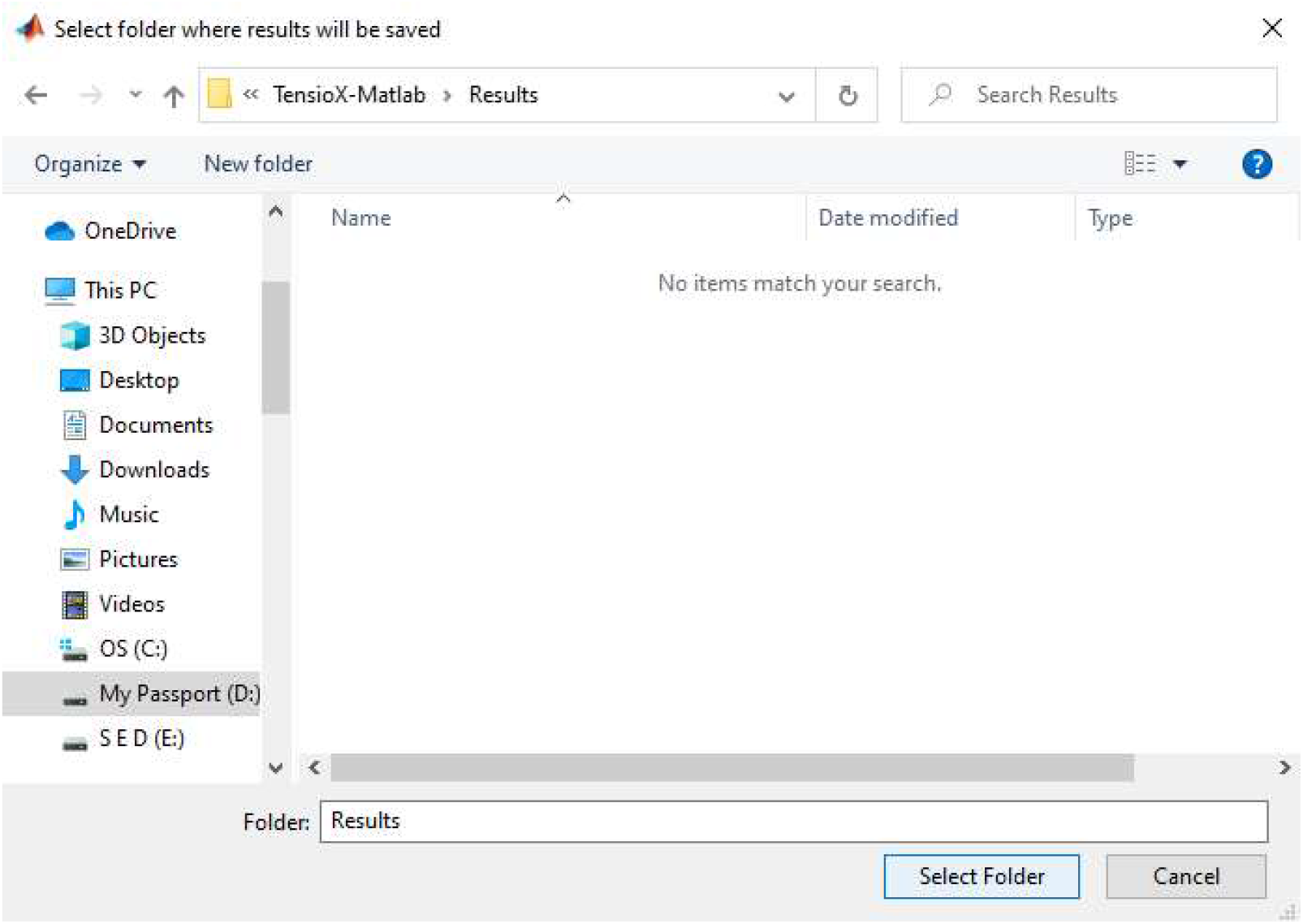
- Spheroid image at *t* = *t*_0_ opens and several windows open asking to select successively the left side, the right side, the apex and the bottom of the spheroid. For each point, press OK then click on the left, right, apex or bottom of the initial spheroid (as shown on the below figure) using the crosshair cursor, then press ENTER. From this, the initial volume of the spheroid is estimated

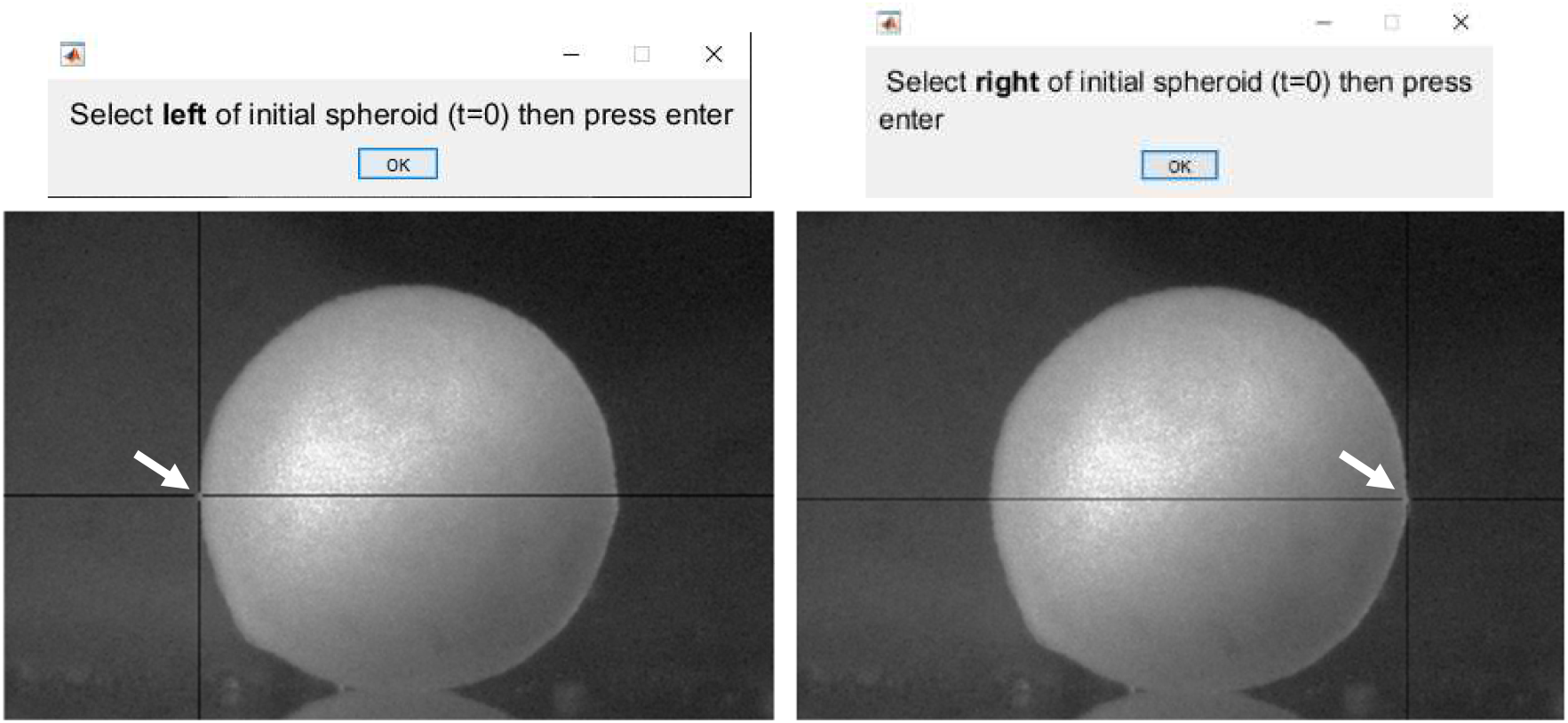
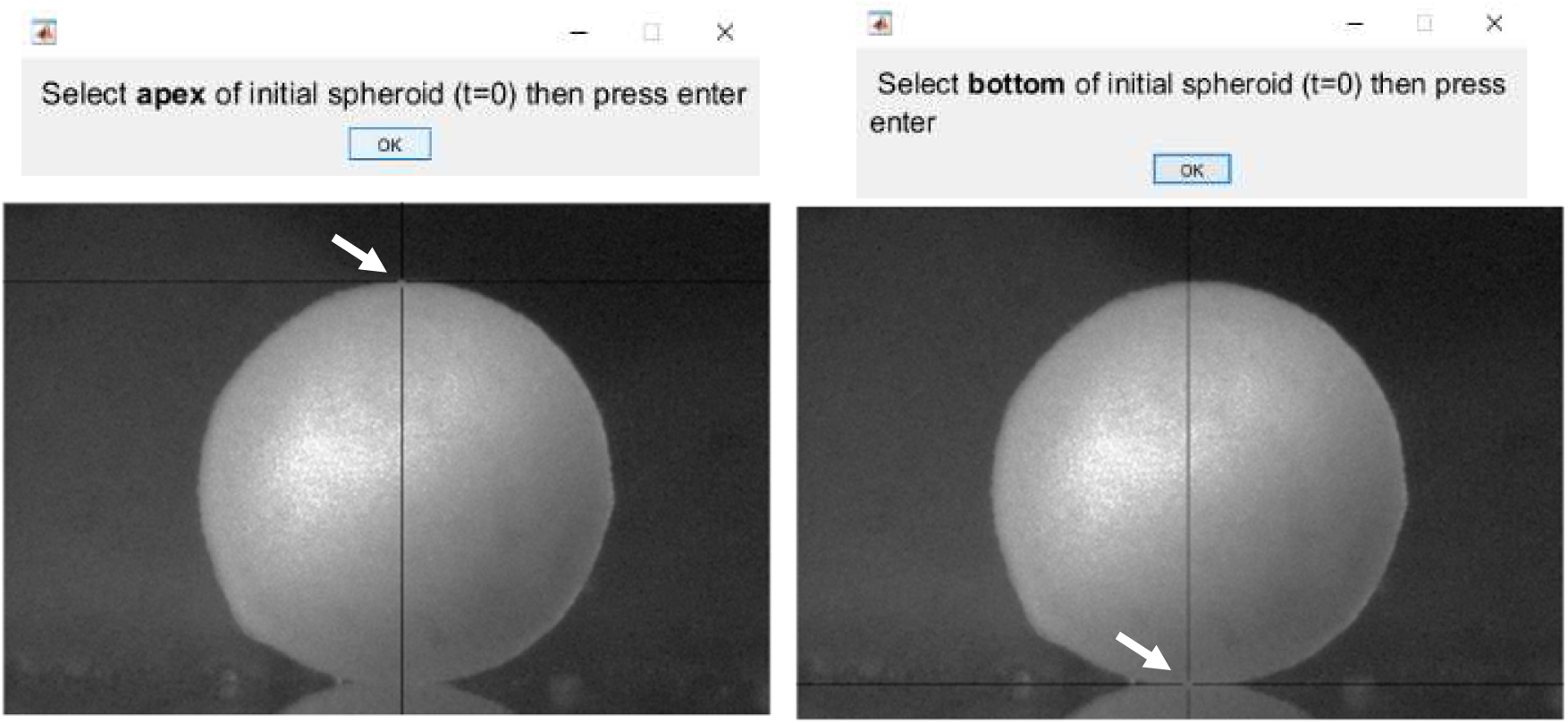
- Spheroid image at *t* = *t*_f_ opens and several windows open asking to select successively the left side, the right side, the apex and the bottom of the spheroid. For each point, press OK then click on the left, right, apex or bottom of the flattened spheroid (as shown on the below figure) using the crosshair cursor, then press ENTER. From this, the theoretical profile is optimized by minimizing the width the height and the volume of the spheroid, to obtain the surface tension of the spheroid.

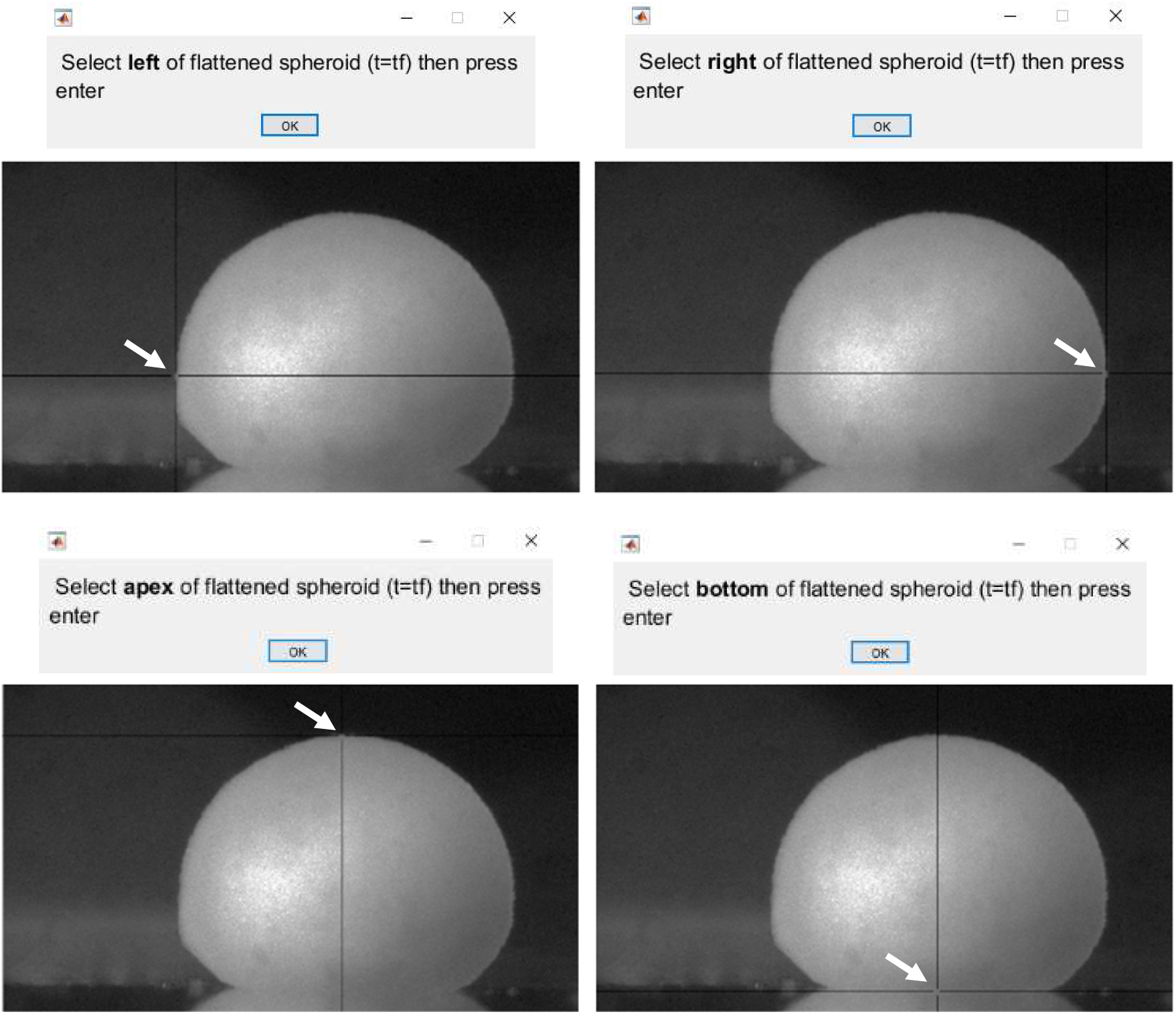
- Finally, two windows open asking to select the right and left limit of the contact area of the flattened spheroid. As previously, for each point, press OK then click on the left or right of the contact area of the flattened spheroid (as shown on the below figure) using the crosshair cursor, then press ENTER.

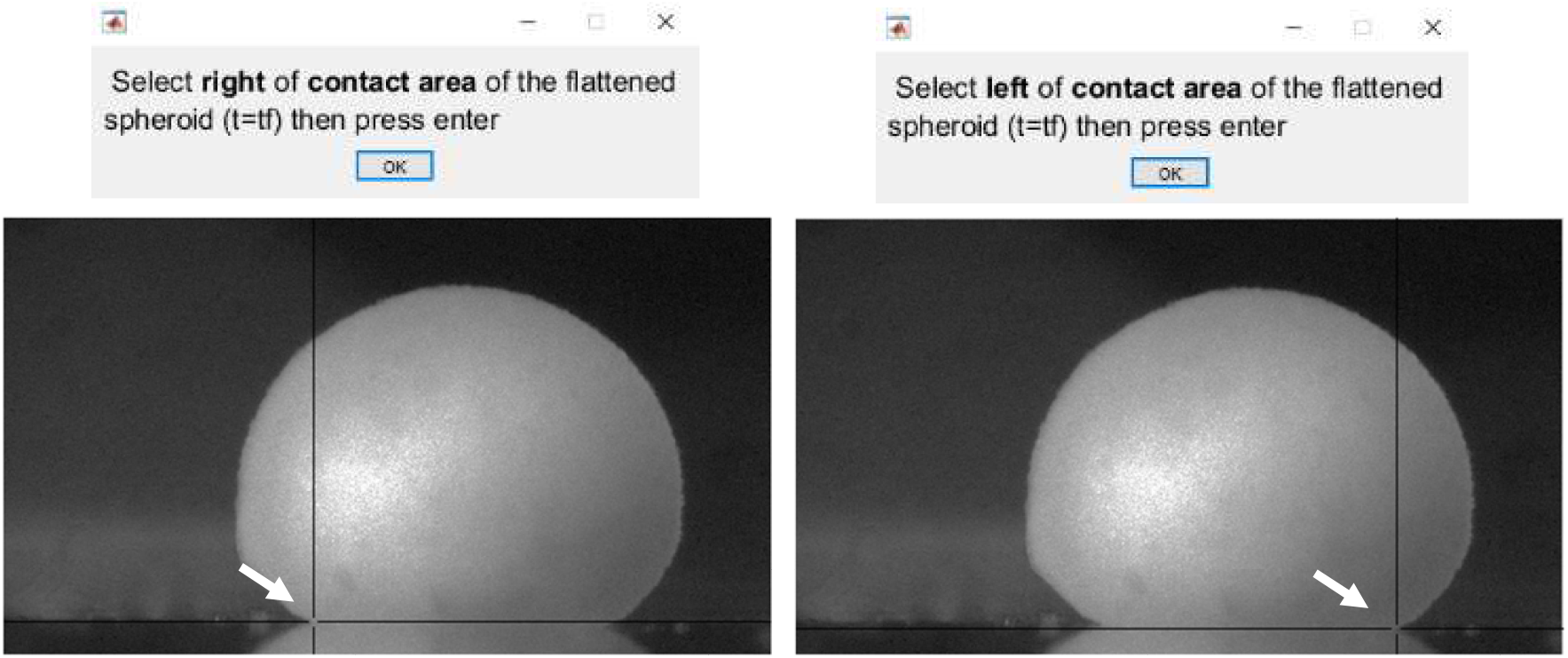
- Enter the image scale in m/pixel, the measured value of the magnetic force per unit of volume in N/m^3^ and an estimated value of gamma in mN/m. If the estimated value of gamma is unknown, leave the default value. Then press OK.

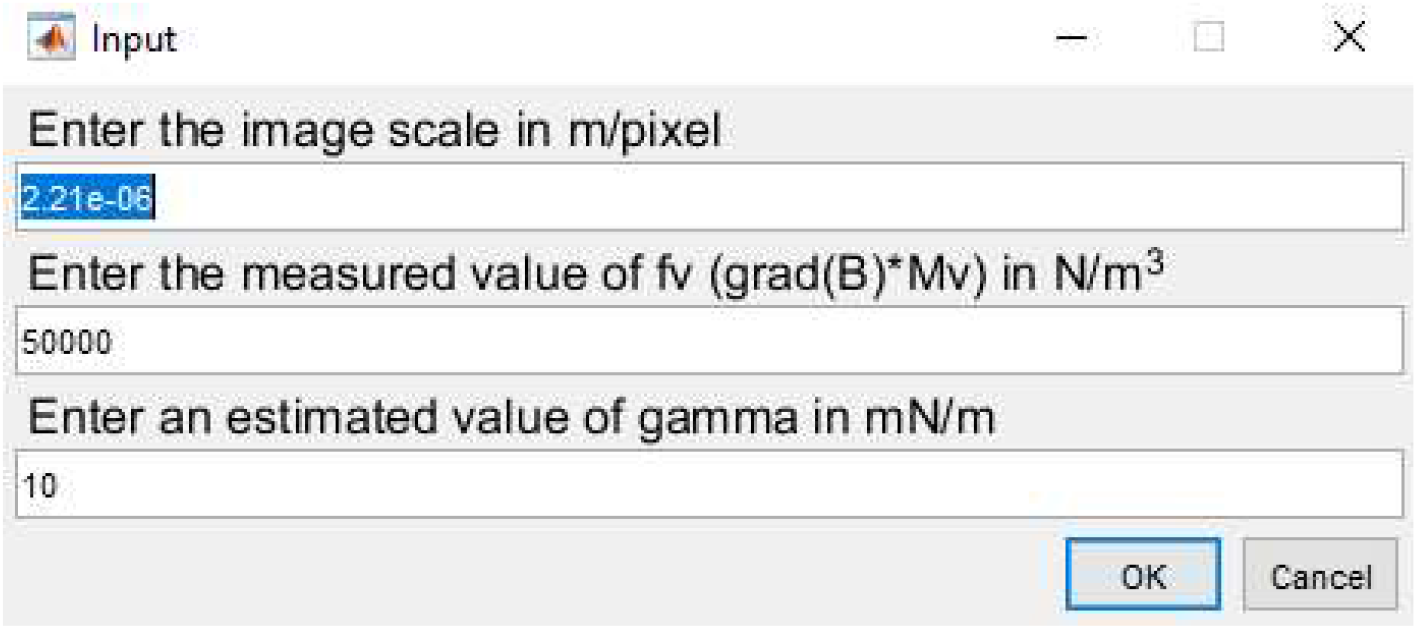
- Two windows open containing the computed surface tension and Young modulus values and an overlay of the flattened spheroid image and the fitted profile for the corresponding surface tension. A good agreement between the experimental and the computed profile (in red) is required to guarantee the reliability of the surface tension results.

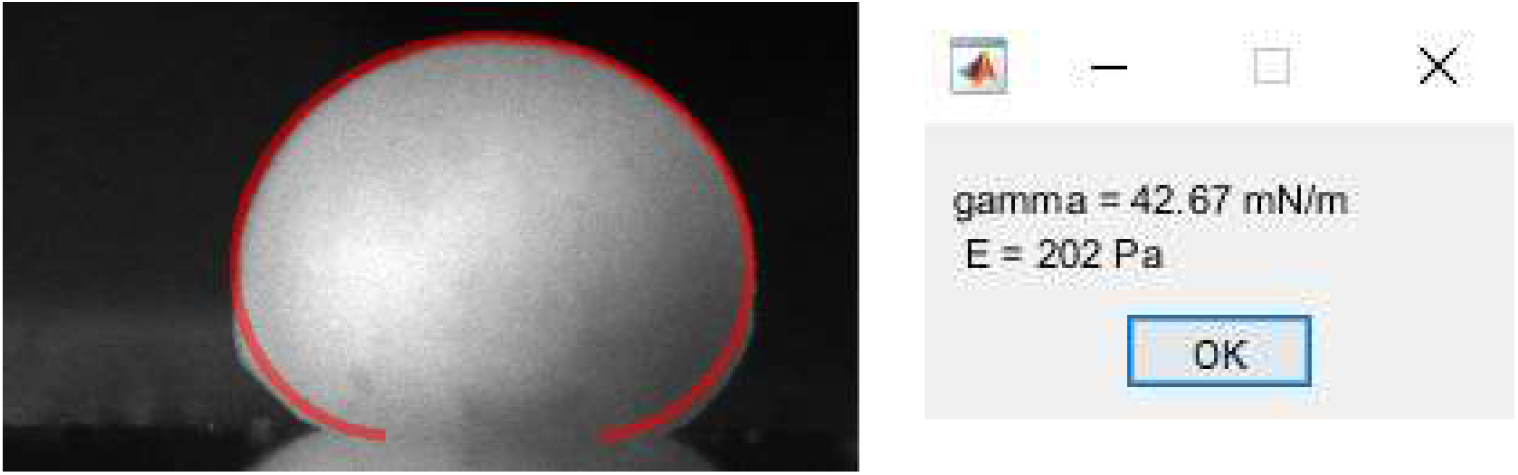
- Both files are saved in the previously selected folder. The computed surface tension and the Young modulus are saved in a results.txt file.

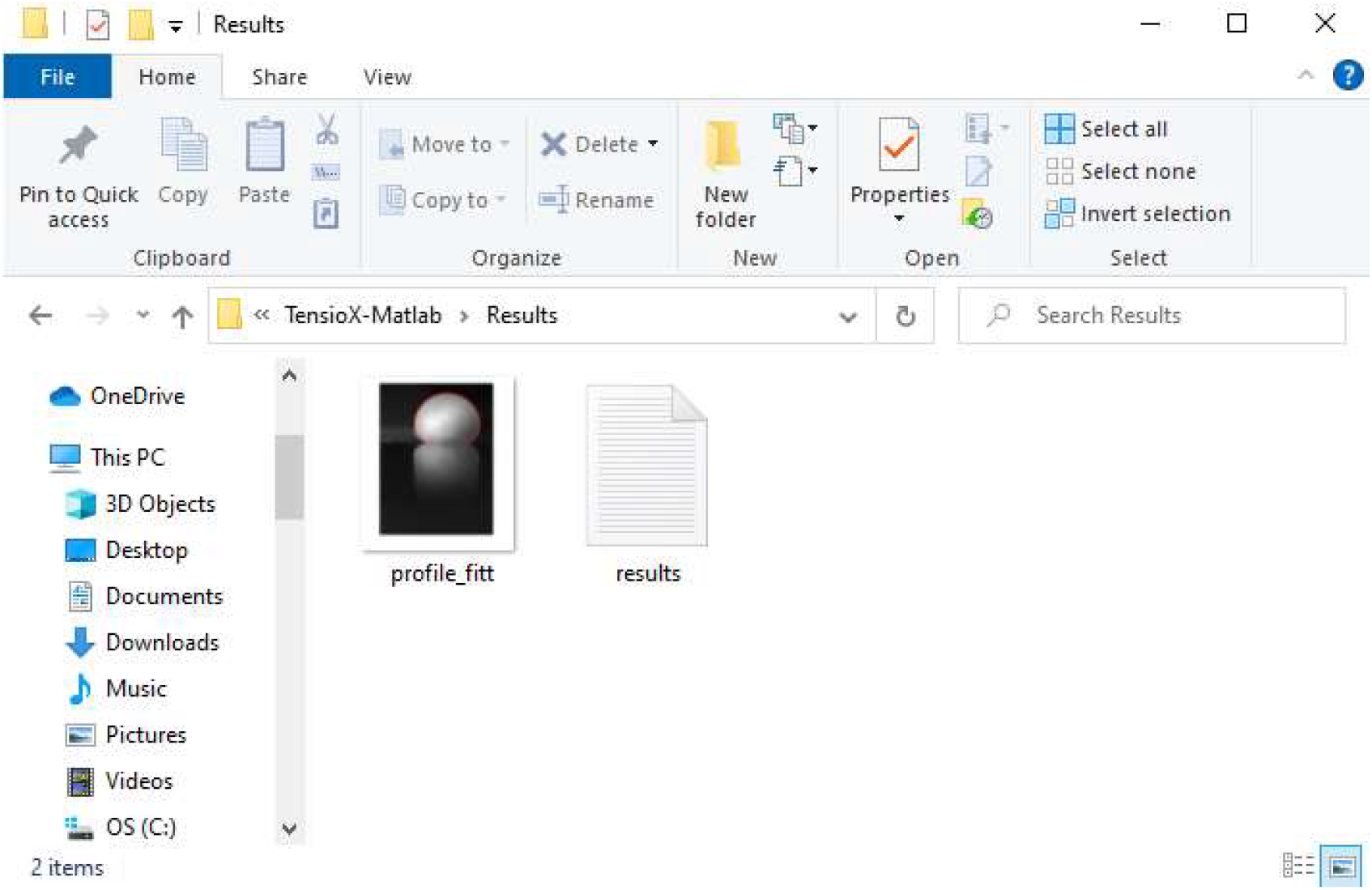

### IV. Detailed description of the MALATBcode

Surface tension and Young modulus are determined as described in [2]. The following paragraphs describe what the MATLABcode contains.

#### 1) Surface tension measurement

Theoretical profiles are obtained by resolving numerically (ode45 function) the classical Laplace equation of capillarity describing the mechanical equilibrium conditions for two homogeneous fluids separated by an interface and in non-wetting conditions. To adjust the numerical profile to the experimental profile, the quadratic error *e* on the height *h*, the width *w* and the volume *V* of the flattened spheroid is minimized with respect to the curvature at apex of the flattened spheroid (*b*) and 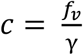 the capillary constant of the system.

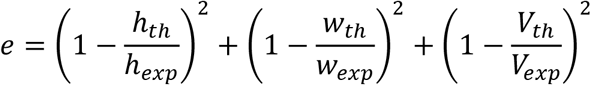

*h*_*exp*_, *w*_*exp*_ are computed from the extracted values on the flattened spheroid image while *V*_*exp*_ is computed from the extracted values on the initial spheroid image (volume of a sphere).

The initial parameters for the minimization are taken such as 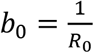 with 2*R*_0_ the average between the width and the height of the initial spheroid and 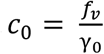 with *f*_v_ and γ_0_ the magnetic force per unit of volume and the estimated surface tension respectively, entered manually. The quadratic error *e* is minimized with the function fminsearch, to obtain the capillary constant of the system from which the surface tension γ is deduced. The final numerical profile and the image of the flattened spheroid are superimposed to check for the reliability of the result.

#### 2) Young modulus measurement

The Young modulus E is computed using Hertz theory for an elastic sphere with an initial radius *R*_0_ which gives 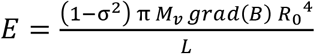 where σ stands for the Poisson ratio (σ = 1/2) and L for the radius of the contact zone computed from the extracted values of the contact area of the flattened spheroid image.

### V. Cautions and remarks

- The initial shape of the spheroid at *t*_0_ has to be spherical to give reliable measurements.
- The substrate on which the spheroid is has to be flat and the camera has to be correctly aligned to provide accurate measurements.
- This application can be adapted to other systems than multicellular aggregates such as any type of viscoelastic fluid or material in non-wetting conditions with respect to the substrate.

